# SCF^Cdc4^ ubiquitin ligase regulates synaptonemal complex formation during meiosis

**DOI:** 10.1101/2020.07.30.228064

**Authors:** Zhihui Zhu, Mohammad Bani Ismail, Miki Shinohara, Akira Shinohara

**Author notes:** College of Science, King Faisal University (Mohammad Bani Ismail). Graduate School of Agriculture, Kindai University (Miki Shinohara). Correspondence: Akira Shinohara, Institute for Protein Research, Osaka University, 3-2 Yamadaoka, Suita, Osaka 565-0871 JAPAN.

## Abstract

Homologous chromosomes pair with each other during meiosis, culminating in the formation of the synaptonemal complex (SC), which is coupled with meiotic recombination. In this study, we showed that a meiosis-specific depletion mutant of a cullin (Cdc53) of the SCF (Skp-Cullin-F-box) ubiquitin ligase, which plays a critical role in cell cycle regulation during mitosis, is deficient in SC formation, but is proficient in the formation of crossovers, indicating uncoupling of meiotic recombination with SC formation in the mutant. Furthermore, the deletion of the *PCH2* gene encoding a meiosis-specific AAA+ ATPase suppresses SC-assembly defect induced by *CDC53* depletion. On the other hand, the *pch2 cdc53* double mutant is defective in meiotic crossover formation, suggesting the SC assembly with unrepaired DSBs. A temperature-sensitive mutant of the *CDC4*, which encodes a F-box protein of the SCF, shows similar meiotic defects to the *CDC53* depletion mutant. These suggest that SCF^Cdc4^, probably SCF^Cdc4^-dependnet protein ubiquitylation, regulates and collaborates with Pch2 in SC assembly and meiotic recombination.

**Summary:** During meiosis, homologous chromosomes pair with each other and form the synaptonemal complex (SC). In this study, components of the SCF (Skp-Cullin-F-box) ubiquitin ligase, Cdc53 and Cdc4, are required for SC formation. A meiosis-specific AAA+ ATPase Pch2 antagonize the functions of Cdc53 and Cdc4 for proper SC assembly.

## Introduction

Meiosis is a specialized cell division, which generates haploid gametes (Marston and Amon, 2004; Petronczki et al., 2003). Upon the entry into meiosis, cells undergo DNA replication followed by two rounds of nuclear division. During meiosis I homologous chromosomes segregate to opposite poles. Crossovers (COs), reciprocal exchanges between homologous chromosomes, are essential for segregation of the chromosomes during meiosis I by providing physical linkages between the chromosomes (Gray and Cohen, 2016; Hunter, 2015).

Meiotic prophase-I exhibits drastic chromosome dynamics and morphogenesis. Homologous loci on two parental chromosomes pair with each other during the prophase-I (Kleckner, 2006; Zickler and Kleckner, 1999). Pairing culminates as synapsis of the homologous chromosomes, manifested by the formation of the synaptonemal complex (SC), a meiosis-specific chromosome structure (Kleckner, 2006; Zickler and Kleckner, 1999). SC contains the central region with polymerized transverse filaments, which is flanked by two homologous chromosomal axes with multiple chromatin loops referred to as a lateral element (LE) (Cahoon and Hawley, 2016; Gao and Colaiacovo, 2018). SC assembly and disassembly are tightly regulated. In leptotene stage, a pair of sister chromatids is folded into a chromosome axis with chromatin loops, called an axial elements (AE). Leptonema is followed by zygonema, in which a short patch of SC is formed between homologous AEs. SC elongation occurs along chromosomes, resulting in the formation of full-length SCs in pachynema, where AEs are referred as to LEs. In some organisms, AE elongation is coupled with SC formation. In other organisms, LE/AE formation proceeds to SC formation. SCs are then disassembled in diplonema, prior to the onset of anaphase-I. Importantly, SC formation is tightly coupled with meiotic recombination in most of organisms including budding yeast and mammals. Mutants defective in meiotic recombination show a defect in SC formation; e.g. the *spo11* and *dmc1* mutant (Baudat et al., 2000; Bishop et al., 1992; Giroux et al., 1989; Romanienko and Camerini-Otero, 2000), which are deficient in the formation of DNA double-strand breaks (DSBs) and strand exchange between homologous DNAs, respectively. On the other hand, in fruit fly and nematode, SC formation is independent of the initiation of meiotic recombination (Dernburg et al., 1998; McKim and Hayashi-Hagihara, 1998).

Synapsis of homologous chromosomes, thus SC formation, initiates at a specific site along chromosomes, which likely corresponds to the site of meiotic recombination. In budding yeast, evolutionally-conserved ZMM (Zip, Msh, Mer)/SIC (Synaptic Initiation Complex) proteins including Zip1, Zip2, Zip3, Msh4, Msh5, Mer3, Spo16, Spo22/Zip4 and Pph3 (Agarwal and Roeder, 2000; Borner et al., 2004; Chua and Roeder, 1998; Hochwagen et al., 2005; Hollingsworth et al., 1995; Nakagawa and Ogawa, 1999; Shinohara et al., 2008; Tsubouchi et al., 2006) promote SC assembly as well as CO formation. ZMMs localize to chromosomes as a large protein ensemble, which is detected by immuno-staining as a focus, for SC assembly through the deposition of Zip1, a yeast transverse filament protein, into arrays in the SC central region (Sym et al., 1993; Sym and Roeder, 1995). Zip1 polymerization is promoted by the action of a complex containing Ecm11 and Gmc2 as a component of the SC central region (Humphryes et al., 2013; Voelkel-Meiman et al., 2013). AEs/LEs contain several meiosis-specific proteins including Red1, Hop1, and Mek1/Mre4 kinase {Hollingsworth, 1990 #81;Leem, 1992 #190;Rockmill, 1988 #23;Rockmill, 1991 #189} as well as a cohesin complex containing a meiosis-specific kleisin Rec8 (Klein et al., 1999). Rec8, Hop1, and Red1 are axis components evolutionally conserved among species and are found as REC8, HORMAD1/2 and SYCP2/3 in mammals, respectively (Eijpe et al., 2003; West et al., 2019; Wojtasz et al., 2009). How AEs or meiotic chromosome axes, which may be independent of SC elongation, are assembled remains largely unknown.

Protein modifications such as ubiquitin and small ubiquitin-like modifier protein (SUMO) regulate various biological processes during both mitosis and meiosis. SUMOlyation is involved in SC formation (Nottke et al., 2017). SUMO is localized to the SC, both the SC central region and the axes, in budding yeast (Cheng et al., 2006; Hooker and Roeder, 2006; Voelkel-Meiman et al., 2013) and both SUMO and ubiquitin are present on the axes and SC central region in mouse spermatocytes (Rao et al., 2017). Budding yeast Ecm11 present in SC central region is SUMOlyated (Humphryes et al., 2013; Voelkel-Meiman et al., 2013)and amounts of SUMOlyated Ecm11 correlates with SC elongation (Leung et al., 2015). In mouse, a SUMO ligase Rnf212 and a ubiquitin ligase Hei10 antagonize with each other for meiotic recombination (Qiao et al., 2014). Moreover, the proteasome is localized on SCs in budding yeast, nematode, and mouse (Ahuja et al., 2017; Rao et al., 2017), suggesting the role of ubiquitin-dependent proteolysis in meiotic chromosome metabolisms.

Two major ubiquitin ligases, the SCF (Skp-Cullin-F-box) and APC/C (Anaphase promoting complex/cyclosome) play an essential role in mitotic cell cycle (Feldman et al., 1997; Skowyra et al., 1997; Yu et al., 1998; Zachariae et al., 1998). In budding yeast meiosis, APC/C with either Cdc20 or Cdh1 promotes the timely transition of metaphase/anaphase I and II (Cooper and Strich, 2011; Pesin and Orr-Weaver, 2008). A meiosis-specific APC/C activator, Ama1, regulates the duration of prophase-I (Okaz et al., 2012), which is negatively controlled by an APC/C subunit, Mnd2 (Oelschlaegel et al., 2005; Penkner et al., 2005).

In budding yeast, a core SCF, which is composed of Rbx1/Hrt1 (RING finger protein), Cdc53 (cullin) and Skp1, binds various F-box adaptor proteins including Cdc4, Grr1, and Met30 (Nakatsukasa et al., 2015; Willems et al., 2004). These F-box proteins determine the substrate-specificity of the complex. SCF with Cdc4, referred to as SCF^Cdc4^, mediates ubiquitylation of G1 cyclin(s) and a Cdk inhibitor, Sic1, at G1/S transition (Koivomagi et al., 2011). The SCF also ubiquitylates Cdc6 essential for the initiation of DNA replication (Perkins et al., 2001). On the other hand, the role of the SCF during prophase-I remains less described. A previous report indicates the role of SCF^Cdc4^ in pre-meiotic DNA replication through Sic1 degradation (Sedgwick et al., 2006).

In this study, we analyzed roles of the SCF ubiquitin ligase in yeast meiosis by characterizing a meiosis-specific depletion mutant of Cdc53 and found that the Cdc53 is indispensable for SC formation and progression into anaphase I. Moreover, a mutant of the *PCH2* gene, which encodes a meiosis-specific AAA+ ATPase (Borner et al., 2008; Chen et al., 2014; San-Segundo and Roeder, 1999) suppresses SC-assembly defect by Cdc53 depletion. A temperature-sensitive *cdc4* mutant also showed similar meiotic defects to the *CDC53* depletion. We propose that SCF^Cdc4^ regulates proper SC assembly by counteracting with the Pch2-dependent negative control on SC assembly.

## Results

### Depletion of Cdc53 induces an arrest at meiosis I

The SCF complex plays a critical role in cell cycle control of mitosis (Willems et al., 2004). However, its role in meiosis is largely unknown, since genes encoding core components of the complex (e.g. Rbx1, Cdc53, Skp1) (Willems et al., 1999; Willems et al., 2004) are essential for vegetative growth of budding yeast, *S. cerevisiae*. To know a meiotic role of the SCF complex in budding yeast, we constructed a strain that depletes an SCF component specifically during meiosis by replacing the promoter of a target gene with the *CLB2* promoter, whose activity is down-regulated during meiosis (Lee and Amon, 2003). In a strain with the *pCLB2-HA-CDC53*, hereafter, *CDC53mn* (meiotic null), we could efficiently reduce a cullin component of the SCF, Cdc53 (Fig. 1 A), whose steady state level did not change much during meiosis (Fig. S1 A). In the *CDC53mn* mutant, the level of Cdc53 began to decrease from 2-h induction of meiosis and little Cdc53 protein was detectable after 4 h. We checked amounts of two known SCF^Cdc4^ substrates, Sic1 and Cdc6. In wild-type cells, the level of Sic1 decreases at 0-2 h incubation with sporulation medium (SPM) (Fig. S1 D), as shown previously (Sedgwick et al., 2006). In the *CDC53mn* mutant, Sic1 was still present at 4 h of meiosis, but disappears at 6 h, suggesting the delay in its degradation. On the other hand, Cdc6, that is very unstable after 4 h in the wild-type meiosis (Perkins et al., 2001), was present at late time points such as 12 h in the mutant (Fig. S1 D). These results indicate that, in the *CDC53mn* mutant, SCF activity is retained in very early meiotic prophase-I, which is enough for triggering premeiotic S phase, but is largely decreased during further incubation.

**Figure 1.**
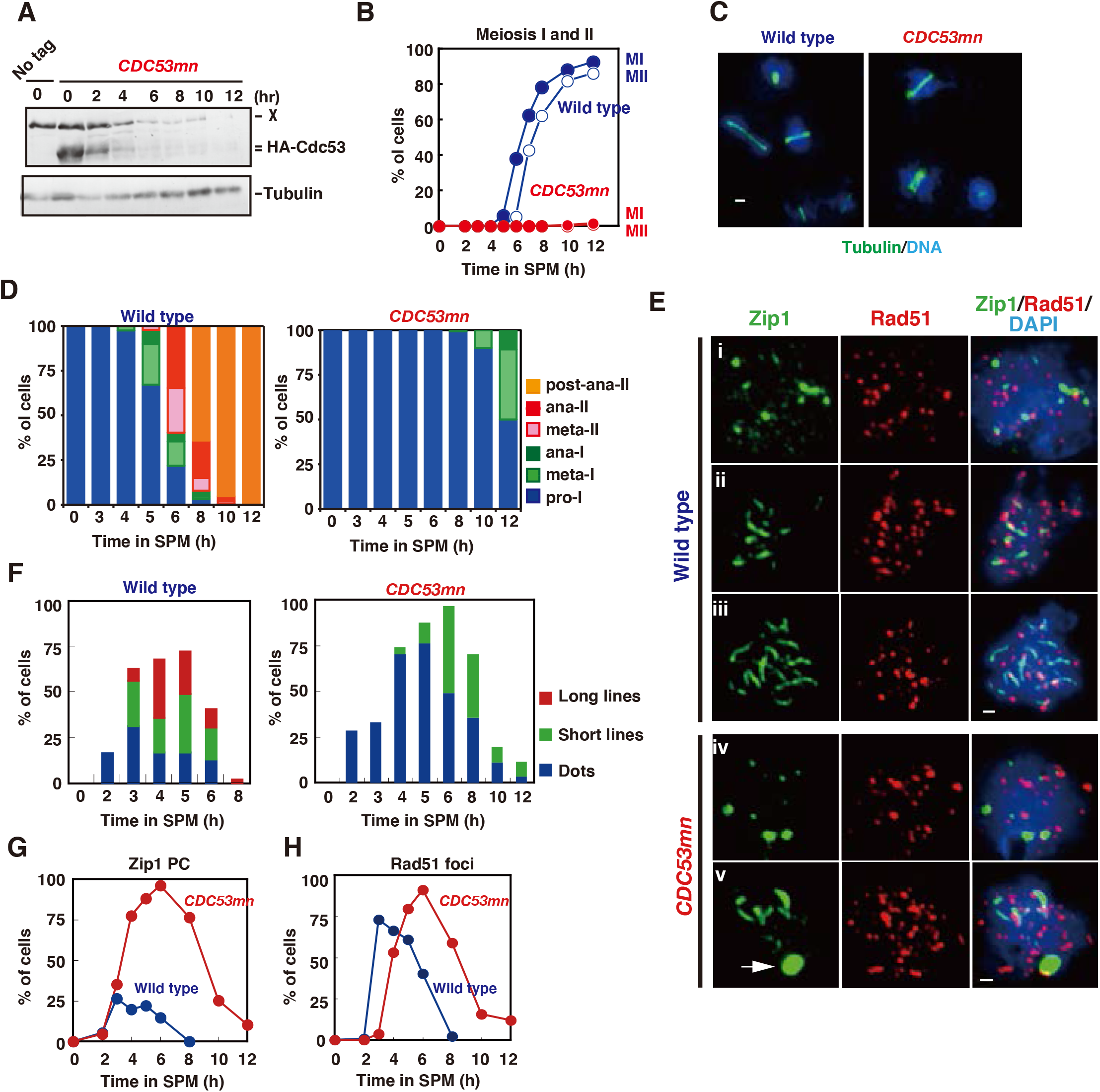
Cdc53-depletion induces abnormal SC. (A) Expression of Cdc53 protein. Lysates obtained from wild type (NKY1551, only at 0 h) and *CDC53mn* (ZHY94) cells at various time points in meiosis were analyzed by western blotting using anti-HA (HA-Cdc53, upper) or anti-tubulin (lower) antibodies. “X” indicates a non-specific band reacted with anti-HA. (B) Meiotic cell cycle progression. Entry into meiosis I and II in the wild-type and *CDC53mn* cells were analyzed by DAPI staining. A number of DAPI bodies in a cell was counted. A cell with 2, 3 and 4 and with 3 and 4 DAPI bodies is defined as a cell which passed meiosis I and meiosis II, respectively. Graph shows the percent of cells that completed MI or MII at the indicated times. More than 200 cells were counted at each time point. Wild type, MI, blue closed circles; Wild type, MII, blue open circles; *CDC53mn* MI, red closed circles; *CDC53mn* MII, red open circles. (C) Tubulin staining in the *CDC53mn* mutant. Whole cells of wild type (5 h) and *CDC53mn* cells (8 h) were fixed and stained with anti-tubulin (green) and DAPI (blue). Representative images are shown. Bar = 2 μm. (D) Classification of tubulin/DAPI staining at each time point of meiosis in wild type (left) and *CDC53mn* mutant (right) cells. Dot, short line and long line tubulin-staining with single DAPI mass are defined as prophase-I, metaphase- and anaphase-I, respectively and shown in different colors. Short and long tubulin-staining are defined as metaphase-II and anaphase-II. At each time point, more than 100 cells were counted. (E) Zip1- and Rad51-staining. Nuclear spreads from wild type and *CDC53mn* mutant were stained with anti-Zip1 (green), anti-Rad51 (red), and DAPI (blue) and categorized. SCs of wild-type cells shown in leptotene (A-I, Class I), zygotene (A-ii, Class II), and pachytene (A-iii, Class III) stages. SCs of *CDC53mn* mutants shown in leptotene (A-iv) and zygotene-like stages (A-v). Representative images are shown. Zygotene-like *CDC53mn* cells contain the polycomplex (PC) as shown by an arrow. Bars = 1 μm. (F) Plots show each class of SC (Wild type, left; *CDC53mn* mutant, right) at indicated times in meiosis. Class I (dots; blue bars), Zip1 dots; Class II (short lines; green bars), partial Zip1 linear; Class III (long lines; red bars), linear Zip1 staining. At each time point, more than 100 cells were counted. (G) The Kinetics of Zip1 polycomplex (PC) formation is shown for each strain. Spreads with Zip1 PC were counted. Wild type, blue; *CDC53mn*, red. (H) Kinetics of Rad51 assembly/disassembly. The number of Rad51-positive cells (with more than 5 foci) was counted at each time point. At each time point, more than 100 cells were counted. Wild type, blue; *CDC53mn* red.

Although the *CDC53mn* mutant shows normal growth during mitosis, the mutant shows various defects during meiosis. The *CDC53mn* mutant cells delayed the onset of meiotic DNA replication by ~2 h compared to the wild-type cells (Fig. S1 B). The delayed Sic1 degradation (Fig. S1 D) might explain the delay in the onset of S-phase in the mutant. DAPI staining showed that Cdc53-depleted cells arrested prior to meiosis I (Fig. 1 B). Aberrant recombination intermediates are known to trigger an arrest at mid-pachytene stage; e.g., *dmc1* mutant (Bishop et al., 1992). However, the arrest in the *CDC53mn* mutant is independent of the recombination. Since the introduction of the *spo11-Y135F* mutation, which abolishes the formation of meiotic DSBs (Bergerat et al., 1997), did not suppress the arrest induced by *CDC53* depletion (Fig. S1 C). We also checked the expression of Cdc5 (Polo-like kinase), which is induced after pachytene exit (Chu and Herskowitz, 1998; Clyne et al., 2003). *CDC53mn* cells expressed Cdc5 from 8 h, 2 h later than the wild type and maintained its expression at late times such as 12 h (Fig. S1 D), showing that the mutant exits the pachytene stage. High levels of Cdc5, whose degradation triggers the exit of meiosis-I, in the mutant support that the mutant does not carry out meiosis I division. Tubulin staining revealed that the mutant delayed the entry into metaphase-I with short spindles (Fig. 1 C). Even at 12 h, only half of the mutant cells contained short metaphase-I or anaphase-I spindles (Fig. 1 D), indicating an arrest at late prophase-I such as metaphase/anaphase-I transition in the *CDC53mn* mutant. These indicate that Cdc53, probably the SCF, plays an essential role in the transition of metaphase-I to anaphase-I, suggesting the presence of a novel regulatory mechanism on the transition. The arrest induced by *CDC53* depletion is similar to that seen in meiosis-specific depletion of the *CDC20*, which encodes an activator or APC/C for the onset of anaphase-I (Lee and Amon, 2003). The SCF may regulate APC/C activity during meiosis I as seen in *Xenopus* oocytes (Nishiyama et al., 2007).

### *CDC53* depletion results in defective SC assembly

We examined the effect of Cdc53-depletion on meiotic prophase-I events such as SC formation by immuno-staining of chromosome spreads. Zip1, a component in SC central region, is widely used as a marker for SC formation (Sym et al., 1993). As a marker for meiotic DSB repair, we co-stained Rad51, a RecA-like recombinase (Bishop, 1994; Shinohara et al., 1992) with Zip1. In wild type, Zip1 first appears as several foci on chromosomes in early meiosis such as leptotene stage (Fig. 1 E i). Then, short lines and later long lines of Zip1 are observed, which correspond with zygotene (Fig. 1 E ii) and pachytene stages (Fig. 1 E iii), respectively. The *CDC53mn* mutant is defective in SC assembly (Fig. 1 E and F). At early time points (e.g., 2-4 h) in the mutant, dotty-staining of Zip1 appeared without any delay compared to wild type. This indicates normal association of Zip1 to chromosomes at early meiosis I. The formation of short lines of Zip1 in the mutant delayed by ~2-3 h compared to that in wild type (Fig. 1 E and F). Even at late time points (6 h), the mutant largely reduced the formation of fully-elongated Zip1-lines (Fig. 1 F), indicating a defect in Zip1 elongation. Indeed, the mutant transiently accumulated an aggregate of Zip1, called “poly-complex (PC)”, which is an indicative for abnormal SC assembly (Sym et al., 1993) (Fig. 1 E and G). Although the mutant transiently accumulated these zygonema-like nuclei with short lines and PCs of Zip1, these Zip1 structures almost dismantled at late times such as 10-12 h in the mutant (Fig. 1 F). The disappearance of Zip1-positive cells is delayed by ~4 h in the mutant relative to wild type, indicating ~2 h longer prophase-I in the *CDC53mn* mutant relative to that in wild type.

### *CDC53* depletion shows little defect in meiotic recombination

During meiotic prophase-I, SC formation is tightly coupled with meiotic recombination (Alani et al., 1990; Bishop et al., 1992; Padmore et al., 1991). Rad51 staining showed that the *CDC53mn* mutant cells delayed the appearance of Rad51 foci on the chromosomes relative to wild-type cells (Fig. 1 E and H), probably due to delayed entry into meiosis (Fig. S1 B). However, kinetics of Rad51-focus staining look similar to those in the wild type, with a delayed peak at 6 h (Fig. 1 H), suggesting a weak defect in DSB repair in the mutant.

To analyze meiotic recombination in the *CDC53*-depletion mutant, we analyzed the repair of meiotic DSBs and formation of crossovers (COs) at a well-characterized recombination hotspot, the *HIS4-LEU2* (Cao et al., 1990) by Southern blotting (Fig. 2 A). In wild type, DSBs appeared at ~3 h, peaked at 4 h and then disappeared after 5 h (Fig. 2 B and C). Consistent with the delay in the onset of meiotic S-phase, the *CDC53mn* mutant delayed DSB appearance by ~2 h relative to wild type. When the delay in the S-phase entry is compensated, kinetics of the appearance of meiotic DSBs in the mutant are similar to those in wild type. On the other hand, there is substantial delay (about one hour) in the disappearance of DSBs in the mutant, indicating a weak defect in meiotic DSB repair. These suggest that the *CDC53mn* mutant is almost proficient in the repair of meiotic DSBs. Indeed, the mutant is proficient in CO formation at this locus. In wild type, COs started to form at 5 h and reached to a maximum level of ~7.5% at 8 h (Fig. 2 D and E). Although the formation of COs in the mutant delayed by ~3 h compared to wild type, final levels of COs in the mutant were indistinguishable from those in wild type (Fig. 2 E). To confirm the normal CO formation at the other locus in the *CDC53mn* mutant, we also analyzed the ectopic CO formation at a recombination hotspot of *URA3-ARG4* (Allers and Lichten, 2001). At this locus, ectopic COs are formed between cassettes at the *leu2* and *his4* loci (Fig. 2 F). The *CDC53mn* mutant showed approximately two-hour delay in the formation of the COs at the locus relative to wild type (Fig. 2 G and H). A final level of CO products in the mutant was similar to those in the wildtype control (Fig. 2 H). In addition, we checked genome-wide DSB repair by examining chromosome bands using pulse-field gel electrophoresis (Fig. S1 E). The *CDC53mn* cells showed smear bands of chromosomes at 4 h, which were caused by DSB formation, and, as wild-type cells, recovered full chromosomal bands from 5 h. This, together with Rad51 focus kinetics (Fig. 1 H), suggests that most of DSBs are repaired under *CDC53*-depletion condition. Furthermore, these results indicate that SC formation is uncoupled with meiotic recombination in the *CDC53mn* mutant. It is likely that full-length SCs are not required for completion of CO formation in the absence of Cdc53.

**Figure 2.**
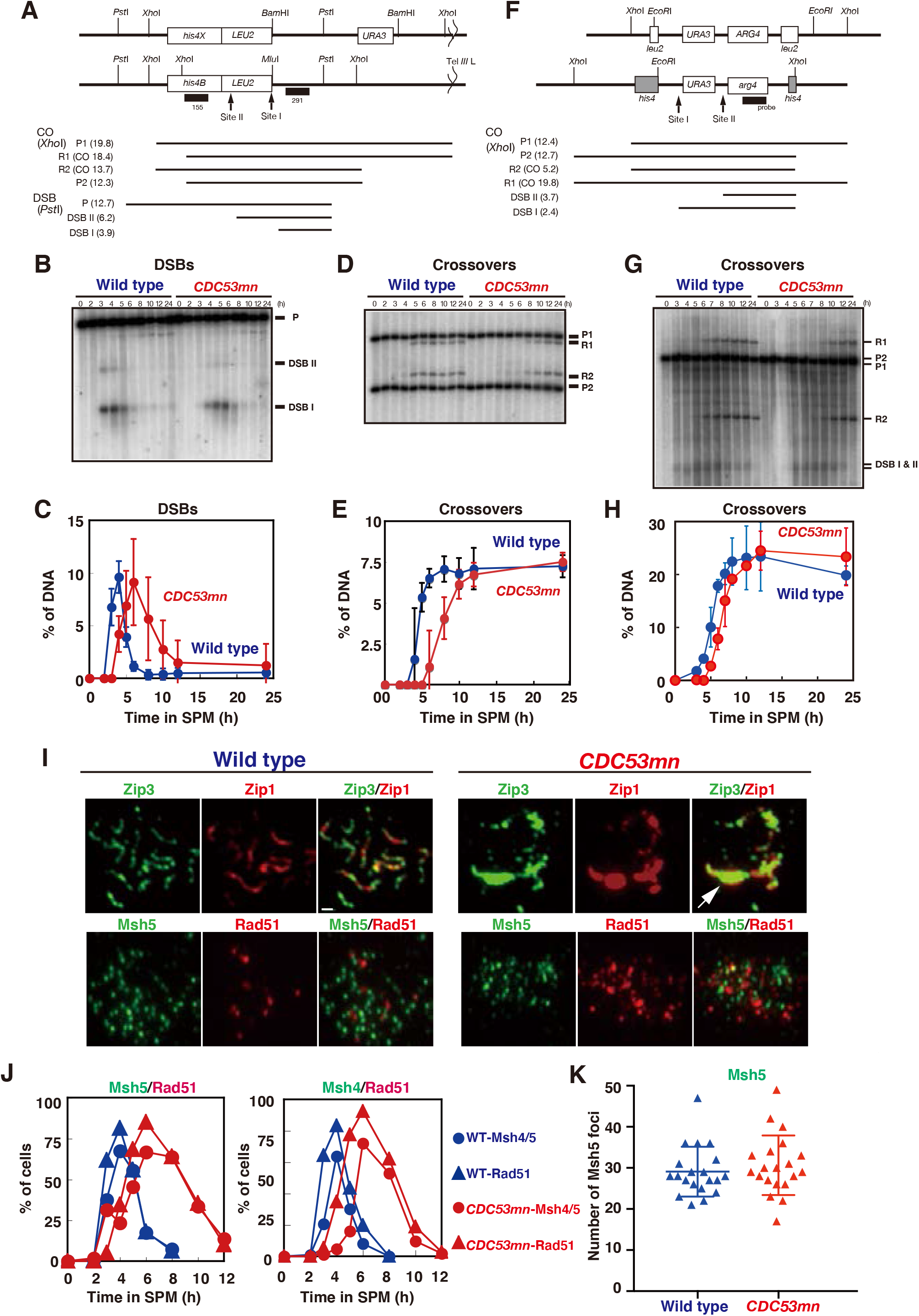
Cdc53-depletion mutant is proficient in meiotic recombination. (A) A schematic diagram of the *HIS4-LEU2* recombination hotspot. Restriction sites for *Pst*I, *Xho*I, *Bam*HI, and *MluI* are shown. Diagnostic fragments for analysis on double-strand break (DSB) and crossover (CO) are shown at the bottom. The size of each fragment (kilo-bases) is presented within parentheses. (B, C) DSB repair at the *HIS4-LEU2* locus were analyzed by Southern blotting (B) and quantified (C). Genomic DNAs were prepared and digested with *Pst*I. Error bars (SD) were obtained from three independent experiments. Wild type, NKY1551, blue circlers; *CDC53mn*, ZHY94, red circles. (D, E) CO formation at the *HIS4-LEU2* locus were analyzed by Southern blotting (D) and quantified (E). Ratios of R1 band to P1 were calculated. Genomic DNAs were digested with *Xho*I. Error bars were obtained from three independent time courses. (F) A schematic diagram of the *URA3-ARG4* recombination hotspot. Restriction sites for *Xho*I, and *EcoRI* are shown. Diagnostic fragments for analysis for parent, DSB and crossover (CO) fragments are shown at the bottom. The size of each fragment (kilo-bases) is presented within parentheses. (G, H) CO formation in *CDC53mn* mutant was verified by Southern blotting. Ectopic CO formation at the *URA3-ARG4* recombination locus were analyzed by Southern blotting (G) and quantified (H). Genomic DNAs were digested with *Xho*I. Error bars were obtained from three independent cultures. Wild type, MJL2442, blue circlers; *CDC53mn*, ASY1202, red circles. (I) Localization of Zip3 and Msh5 in the *CDC53mn* mutant. Chromosome spreads from wild type (4 h, NKY1551) and *CDC53mn* mutant (8 h, ZHY94) were stained with anti-Zip3 or anti-Msh5 antibodies (green) together with anti-Zip1 (red). Representative images are shown. Bar = 1 μm. (J) Assembly of Msh4-Msh5 in the *CDC53mn* mutant. The percent of cells positive for Msh4, Msh5, or Rad51 foci (more than 5 foci per nucleus) were counted at each time point. At least 100 nuclei were counted at each time point. Wild type, blue triangles and circles; *CDC53mn* mutant, red triangles and circles. Triangles and circles are for Rad51 and Msh4 (right) or Msh5 (left), respectively. (K) The number of Msh5 foci per a spread was counted at 4 h in wild-type (NKY1551) at 6 h in the *CDC53mn* mutant (ZHY94). 20 spreads were counted. Mean and SD are shown on the plot.

### *CDC53* depletion results in altered assembly of some ZMM proteins

We examined the localization of ZMM proteins (Zip3, Zip2, Mer3, Spo22/Zip4, Msh4, and Msh5), which promote SC assembly and CO formation. As reported previously (Agarwal and Roeder, 2000; Chua and Roeder, 1998; Shinohara et al., 2008; Tsubouchi et al., 2006), on wild-type chromosomes these ZMM proteins show punctate staining from leptotene to pachytene stages (Figs. 2 I and S2 A). We found that the staining of Zip3, Zip2, Mer3, and Spo22/Zip4 were altered in the *CDC53mn* mutant (Figs. 2 I, top and S2 A). With reduction of focus staining, the mutant accumulated PCs of Zip3, Zip2, Mer3 and Spo22/Zip4, which almost colocalize with Zip1 PCs (Figs. 2 I and S2 A). These indicate defective assembly of ZMM proteins in the *CDC53mn* mutant.

Different from foci of Zip2, Zip3, Mer3 and Spo22/Zip4 proteins, foci of Msh4 and Msh5, which form a hetero-dimeric MutSγ complex (Hollingsworth et al., 1995; Novak et al., 2001), look normal in the *CDC53mn* mutant compared to those in wild type (Figs. 2 I and S2 A). In the mutant, Msh4 and Msh5 foci appeared with a 2-h delay, and disappeared with delay (4 h) relative to wild type (Fig. 2 J). As in wild type, the kinetics of Msh4 and Msh5 are similar to those of Rad51 in the mutant (Figs 1 H and 2 J). Notably, the *CDC53mn* mutant does not form any PCs containing Msh4 or Msh5 (Fig. 2 I and S2 A), different from other SC-defective mutants such as *zmm* (Shinohara et al., 2015; Shinohara et al., 2008). We also counted a steady-state number of Msh5 foci in the *CDC53mn* mutant. An average number of bright Msh5 foci at 6 h in the mutant (Fig. 2 K) was 30.7±7.3 (n=20) which is indistinguishable from the number at 4 h in wild type (29.1±6.1; Mann-Whitney *U*-test, *P*=0. 34). Normal assembly and disassembly of Msh4 and Msh5 could explain the proficiency of meiotic CO formation in the mutant (Fig. 2 E and H). This result supports the idea that, among ZMMs, Msh4-Msh5 is a key for crossover formation, whose functions are distinct from other ZMMs (Pyatnitskaya et al., 2019; Shinohara et al., 2008). Moreover, Cdc53 plays a role in proper assembly of subsets of ZMM proteins other than Msh4-Msh5 for SC assembly.

### Cdc53 and Zip3 distinctly work in SC formation and recombination

To know the relationship between Cdc53 (ubiquitin ligase) and Zip3 (SUMO ligase), both of which is involved in SC assembly, we characterized the *CDC53mn zip3* double mutant. Like the *CDC53mn* single mutant, the *CDC53mn zip3* double mutant shows an arrest during meiosis (Fig. 3 A), which is different from the *zip3* mutant with delayed progression of the meiotic prophase (Agarwal and Roeder, 2000; Borner et al., 2004). While both of the *CDC53mn* and *zip3* single mutants can repair meiotic DSBs with substantial delay, *CDC53mn zip3* double mutant accumulated unrepaired DSBs at the *HIS4-LEU2* hotspot with hyper-resection at late time points (Fig. 3 B and C). The inability of the double mutant to repair the DSBs was confirmed by the accumulation of Rad51 foci at late time points such as 12 h (Fig. 3 D), when the disappearance of Rad51 foci is seen in both *CDC53mn* and *zip3* single mutants. Moreover, *CDC53mn zip3* double mutant formed lower levels of COs compared to the *zip3* single mutant (Fig. 3 E and F). Different from *CDC53mn* single mutant, the double mutant did not express Cdc5 as a marker for pachytene exit (Chu and Herskowitz, 1998) (Fig. S2 B), indicating an arrest induced by recombination checkpoint response to unrepaired DSBs. These results suggest that Cdc53 plays a role for efficient repair of meiotic DSBs, thus CO formation, in the absence of Zip3.

**Figure 3.**
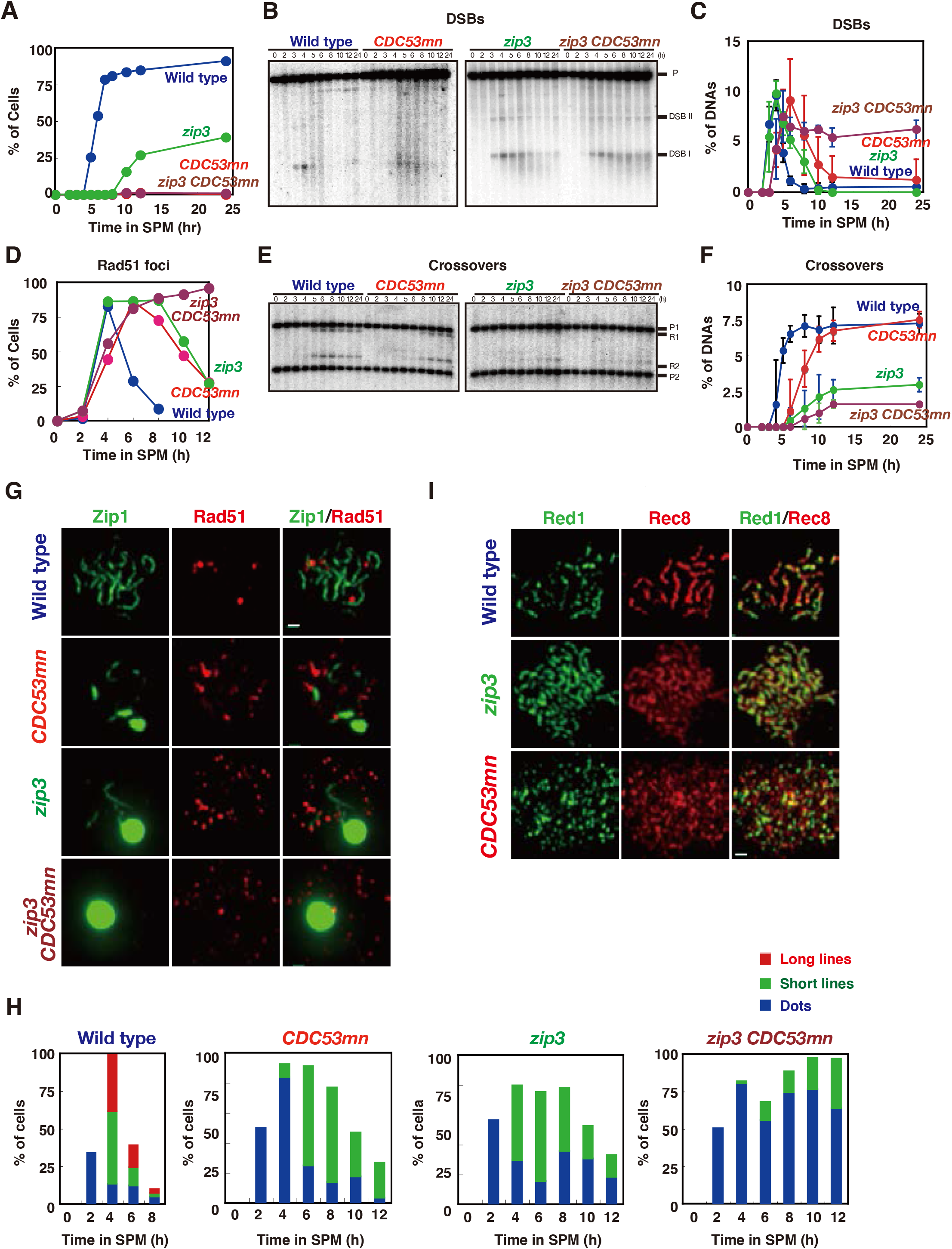
Cdc53 and Zip3 work independently in SC formation and meiotic recombination. (A) Cell cycle progression of various mutants. Entry into meiosis I in the wild type (NKY1551, blue), *CDC53mn* (ZHY94, red), *zip3* (MSY2889, green), *zip3 CDC53mn* mutant (ZHY259, brown) cells were analyzed by DAPI staining/counting as described in Fig. 1B. (B, C) DSB repair at the *HIS4-LEU2* locus in various strains was analyzed as described above. Blots are in (B) and quantification is shown in (C). Error bars were obtained from three independent cultures. (D) Rad51 staining in various mutants was analyzed as described above. Typical staining patterns of each mutant are shown in (G). At least 100 spreads were counted at each time point. Wild type, blue; *CDC53mn*, red; *zip3*, green; *zip3 CDC53mn*, brown. (E, F) CO formation in various strains was analyzed at the *HIS4-LEU2* locus as described above. Blots (E) and quantification (F) are shown. Error bars were obtained from three independent experiments. (G, H) Chromosome spreads from wild-type, *CDC53mn, zip3, zip3 CDC53mn* cells were stained with anti-Zip1 (green) as well as anti-Rad51 (red) and the staining pattern for Zip1 was classified into classes and plotted (H) as shown in Fig. 1F. Bar = 1 μm (I) Chromosome axis formation was analyzed by staining chromosome spreads from various strains with anti-Red1 (green) and anti-Rec8 (red). Wild type, 4 h; *CDC53mn*, 8 h; *zip3*, 8 h; *zip3 CDC53mn*, 8 h. Bar = 1 μm.

SC formation in the *CDC53mn zip3* double mutant was analyzed by Zip1 staining. On spreads of both the *CDC53mn* and *zip3* single mutants, short lines of Zip1 were often formed (Fig. 3 G and H). On the other hand, the *CDC53mn zip3* double mutant reduces the formation of short Zip1 lines compared to either the single mutant (Fig. 3 H). Different from the either single mutants, which disassemble abnormal SCs at late times, the double mutant did not disassemble SCs (Fig. 3 H), which is probably due to pachytene arrest (Fig. S2 B). These suggest that Cdc53 and Zip3 distinctly work for SC assembly as well as CO formation.

### *CDC53mn* mutant forms altered chromosome axis

Defective SC assembly in the *CDC53mn* cells is due to either the assembly of SC *per se* or rapid turnover (precocious disassembly) of fully-elongated SCs in the mutant. To distinguish these possibilities, we introduced an *ndt80* mutation, which blocks the disassembly of the SC by inducing mid-pachytene arrest (Fig. S3A)(Xu et al., 1995). As shown previously (Xu et al., 1995), the *ndt80* single mutant accumulated full length SCs (Fig. S3 B-D). The *ndt80* mutation weakly suppresses SC-elongation defect in the *CDC53mn* mutant only at late times; e.g. 10 h (Fig. S3 D). However, about 20% of the *CDC53mn ndt80* double mutant showed long Zip1 lines at late times, while few long Zip1 lines were transiently formed upon Cdc53 depletion, indicating that the SC assembly defect in the *CDC53mn* mutant is not caused by precocious SC disassembly. Based on above results, we concluded that Cdc53 is necessary for efficient SC assembly.

We confirmed defective SC formation in the mutant by analyzing the localization of chromosomal axis proteins, Hop1 (Fig. 4 A). In the wild-type cells, Hop1 initially binds to unsynapsed chromosomes as multiple foci/lines and then later large fractions of Hop1 a dissociate from synapsed chromosomes (Fig. 4 A), as shown previously (Smith and Roeder, 1997). The *CDC53mn* mutant accumulated Hop1 on chromosomes as multiple foci (Fig. 4 A and B). Even at late times (6 and 8 h), the multiple Hop1 foci/lines persisted on the chromosomes and disappeared at late times. The appearance and disappearance of Hop1 delayed in the mutant by ~2 h and ~4 h delay, respectively, relative to wild type (Fig. 4 B).

**Figure 4.**
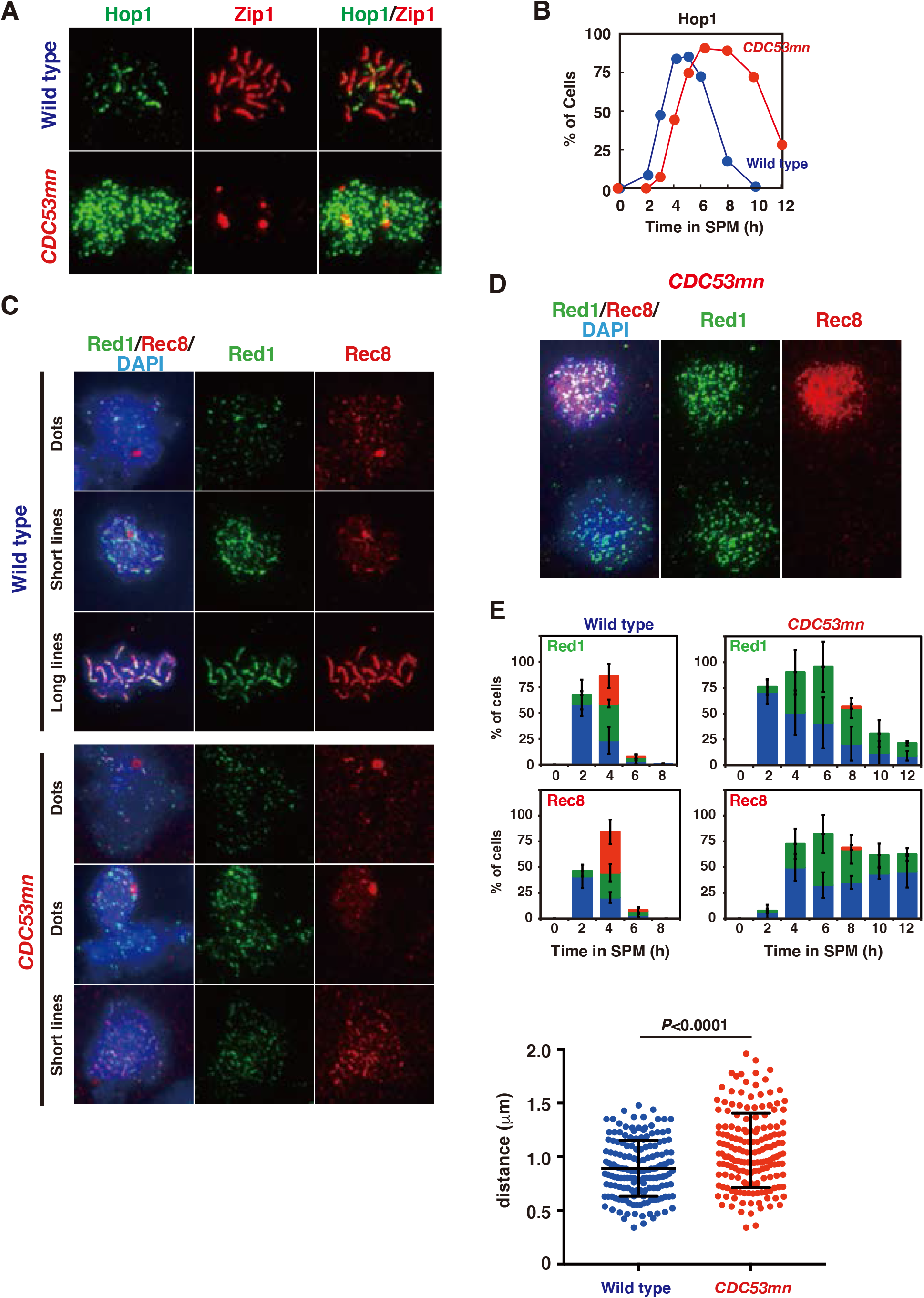
Cdc53 depletion induces altered axis formation. (A) Hop1 staining in wild type (NKY1551) and the *CDC53mn* mutant (ZHY94). Chromosome spreads in each strain were stained with anti-Hop1 (green) and anti-Zip1 (red). For the wild type, pachytene chromosomes with long Zip1 lines are shown with a few Hop1 foci. Representative images are shown. (B) Kinetics of Hop1 assembly/disassembly. The number of Hop1-positive cells was counted at each time point. More than 50 spreads were counted. Wild type, blue circles; *CDC53mn*, red circles. (C, D) Nuclear spreads from wild type and *CDC53mn* mutant were stained with anti-Red1 (green), anti-Rec8 (red), and DAPI (blue) and categorized. SCs of wild-type cells shown in dots, short lines, and long lines. SCs in the *CDC53mn* mutants shown in dots and short lines. Staining of Rec8/Red1 in two adjacent chromosomes spreads of the mutant at 3 h is shown in (D). Top is positive for both Rec8 and Red1. The bottom is positive for Red1, but negative for Rec8. Bars = 1 μm. (E) Kinetics of each class of Red1 (upper two graphs) and Rec8 (lower two graphs) in wild type and *CDC53mn* mutant at indicated times during meiosis. Dots (blue bars), short lines (green bars), long lines (red bars). At each time point, more than 50 spreads were counted. Error bars are SD (*n*=4). (F) Chromosome compaction was measured using cells with both GFP-marked CenIV and TelIV. Distance between two GFP signals was measured using NIH Image J and plotted. Wild type, ZHY749 (blue); *CDC53mn*, ZHY 750 (red). Means (*n*=166) and S.D. are shown. *P* value was calculated with Mann Whitney U test.

We also examined the localization of other axis components, Rec8 and Red1 (Klein et al., 1999; Smith and Roeder, 1997). Wild-type cells show dot/short line staining of Red1 and Rec8 in early prophase-I. In pachytene stage, when synapsis is almost completed, both proteins show beads-in-line staining (Fig. 4 C), which is different from Hop1. In wild-type spreads, both Rec8 and Red1 showed dots at 2 h and short and long lines with beads-in-line staining at 4 h (Fig. 4 C). In wild type, Red1 and Rec8 disappeared at 6 h (Fig. 4 E). On the other hand, the *CDC53mn* mutant formed few Red1/Rec8 long beads-in-lines at any times during prophase-I (Fig. 4 C and E), consistent with SC defects in the mutant. Moreover, in the *CDC53mn* mutant, Red1 dots appeared from 2 h, short-line staining peaked at 6 h and, from 8 h, Red1 signals gradually decreased during further incubation (Fig. 4 E). Contrary to Red1, few Rec8 dot-positive spreads was observed at 2 h in the mutant. Short lines of Rec8 appeared from 4 h and peaked at 6 h and some fractions of Rec8 short-lines started to disappear from 8 h, probably due to the cleavage-independent cohesin release (Challa et al., 2019). Rec8 dots persisted on chromosomes during further incubation as a result of metaphase-I arrest; no cleavage of Rec8. This result indicated the uncoordinated loading of axis proteins Red1 and Rec8/Hop1 in early meiosis in the *CDC53mn* mutant. Indeed, the mutant showed spreads at early time points such as 3 h, which are positive for Red1 but negative for Rec8 (Fig. 4 D). This staining was rarely seen in early wild-type spreads (Fig. 4 C). Rather than delayed loading of Hop1/Rec8, the mutant exhibited precocious loading of Red1 (and Zip1; Fig. 1 F), suggesting un-coordinated loading of chromosomal proteins. These imply that the *CDC53mn* mutant assembles an altered structure of meiotic chromosome axis.

We also analyzed chromosome axis structure by deconvolution analysis of DAPI-stained chromosomes. In wild type, we often see two pairs of DAPI-stained lines are co-aligned with each other (Fig. 5 A). Like the *zip3* mutant (Agarwal and Roeder, 2000), the *CDC53mn* mutant formed few DAPI-dense linear structures. Thus, these suggest that Cdc53 might promote proper formation of chromosome axes. To probe chromosome axis structure, we compared Rec8 staining in the *CDC53mn* mutant with those in the *zip3* and *gmc2* mutants, both of which are defective in SC formation. In the *zip3* mutant, which shows zippering-defect of SCs (Agarwal and Roeder, 2000), beads-on-line staining of both Rec8 and Red1 is observed at late time points, suggesting normal assembly of AE or chromosome axis in the *zip3* mutant. On the other hand, the *CDC53mn* mutant shows more dotty staining of Rec8 than the *zip3* (Fig. 3 I). The *gmc2* mutant is also defective in SC elongation, but retains ability to forms COs (Humphryes et al., 2013), which is similar to the *CDC53mn* single mutant. Unlike the *CDC53mn* mutant, the *gmc2* mutant also exhibited elongated Rec8 lines (Fig. S3 E). This supports the idea that the *CDC53mn* mutant form altered chromosome axis, which is different from other SC-defective mutants.

**Figure 5.**
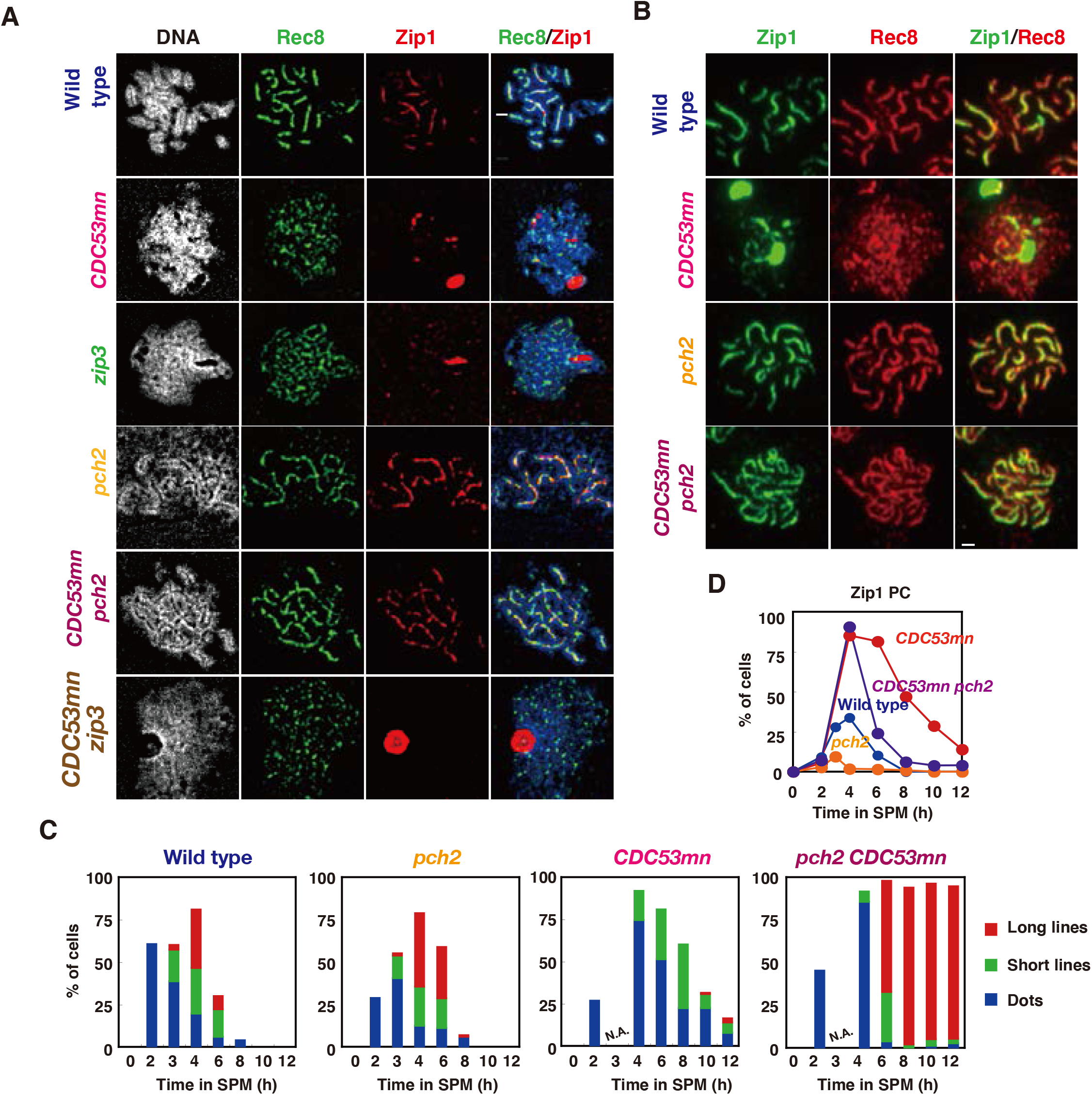
The *pch2* mutation suppresses SC defects induced by Cdc53 depletion. (A) Chromosome spreads from various strains were stained with anti-Zip1 (red) and anti-Rec8 (green) as well as DAPI (white) and images were captured using DeltaVision eipi-fluorescent microscope and deconvoluted as described in Materials and Methods. Representative images are shown. Wild type, NKY1551; *CDC53mn*, ZHY94; *zip3*, MSY2889; *pch2*, ZHY350; *pch2 CDC53mn*, ZHY351; *zip3 CDC53mn*, ZHY259. Bar = 1 μm. (B) Chromosome spreads from various strains were stained with anti-Rec8 (green) and anti-Zip1 (red). Wild type, 4 h; *CDC53mn*, 8 h; *pch2*, 6h; *pch2 CDC53mn*, 8 h. Bar = 1 μm. (C) Zip1-staining in each strain were classified and plotted at each time point. More than 100 nuclei were counted as shown in Figure 1(F). Class I (blue bars), Zip1 dots; Class II (green bars), partial Zip1 linear staining; Class III (red bars), linear Zip1 staining. N.A. indicates “unavailable”. (D) Cells containing Zip1 polycomplexes (PCs) were counted at each time point and plotted. Wild type, blue; *CDC53mn*, red; *pch2*, orange; *pch2 CDC53mn*, purple.

During meiotic prophase-I, chromosome axes are compacted compared to that in pre-meiotic cells (Challa et al., 2019; Challa et al., 2016). We measured the distance between CenIV and TelIV loci marked with GFP in an intact cell (Fig. S2, Fig. 4 F). The distance of the two loci in wild-type cells at 4 h was 0.89 ± 0.26 μm (*n*=166) while that at 5 h in the *CDC53mn* mutant cells was 1.06 ± 0.35 μm (*n*=166, Mann Whitney, *P* value<0.0001). This showed that the *CDC53mn* mutant is defective in chromosome compaction during meiosis.

### The *pch2* deletion largely suppresses SC defect by Cdc53 depletion

As a SCF component, Cdc53 facilitates the degradation of a target protein which functions as a negative regulator for biological processes; e.g. Sic1. We hypothesized that Cdc53 may relieve the negative regulation on SC formation and looked for a mutant which rescues a defect in the *CDC53mn* mutant. We found that the deletion of the *PCH2* gene largely suppresses SC defects induced by Cdc53 depletion (Fig. 5 A and B). The *pch2* mutant was originally isolated by one which alleviates meiotic prophase-arrest induced by the *zip1* mutation (San-Segundo and Roeder, 1999). Pch2 is a conserved AAA+ (ATPase Associated with various cellular Activity) ATPase protein (Wu and Burgess, 2006) involved in remodeling of chromosome axes by modulating Hop1 (Borner et al., 2008; Chen et al., 2014) as well as in the pachytene checkpoint and regulation of DSB formation (Wu and Burgess, 2006). As described previously (Borner et al., 2008; San-Segundo and Roeder, 1999), the *pch2* single mutant shows more continuous Zip1 lines than wild-type (Fig. 5, A and B)(Borner et al., 2008). Different from the *CDC53mn* single mutant, the *CDC53mn* mutant with the *pch2* deletion did form uniformly-stained long Zip1-lines like the *pch2* single mutant (Fig. 5, A and B). Staining of Rec8 revealed similar Rec8 lines in the *CDC53mn pch2* double mutant like the *pch2* single mutant (Fig. 5, B), suggesting that the *pch2* suppresses the SC defect in the *CDC53mn* mutant.

We checked whether SCs in the double mutant are formed between homologous or non-homologous chromosomes. Pairing of a centromere locus was tested using CenXV-GFP (Fig. S4D). Wild-type spreads harboring one spot peaked at 5 h with 30% (n=50), probably due to transient nature of pairing at the locus. The *CDC53mn* showed similar frequency of 28%, (at 6 h) supporting that the mutant is proficient in the pairing. The *CDC53mn pch2* double mutant accumulated spreads with one spot with a frequency of 74% at 8 h, indicating normal pairing of CenXV-GFP in the double mutant. These indicate that SCs in the double mutant are formed between homologous chromosomes.

The *CDC53mn pch2* double mutant initially accumulated Zip1 PCs with defective SC assembly at earlier times as seen in the *CDC53mn* single mutant, but, during further incubation, the PCs disappeared and concomitantly full-length SCs appeared in the double mutant (Fig. 5, C and D). At 8 h, most of the nuclei of the double mutant contained full SCs without any PCs. These suggest that suppression of SC defects by the *pch2* mutation is not due to suppression of early defects conferred by Cdc53 depletion. Consistent with this, neither delayed onset of S-phase nor delayed degradation of Sic1 and Cdc6 in *CDC53mn* cells was suppressed by the *pch2* (Fig. S4 A and B). The effect of the *pch2* deletion on the suppression of SC-defects is very specific to Cdc53-deficiency in SC assembly, since the *pch2* deletion mutation does not suppress a SC-defect induced by the *zmm* mutation (San-Segundo and Roeder, 1999).

Although the *CDC53mn pch2* double mutant looks to form normal SCs, the mutant arrested at the pachytene stage with full length SCs and did not disassemble SCs or exit this stage (Fig. 5 C and 6 A). We next checked meiotic recombination. The *CDC53mn pch2* double mutant could not repair meiotic DSBs at *HIS4-LEU2* locus and accumulated more processed DSB ends (Fig. 6 B and C), which was accompanied with the accumulation of Rad51 foci on spreads (Fig. 6 D). Consistent with DSB repair defect, the *CDC53mn pch2* double mutant largely decreased CO levels compared to the wild-type strain (~1/5 of the wild type level; Fig. 6 E and F). On the other hand, the *CDC53mn* (Fig. 3) and *pch2* single (Borner et al., 2008; Hochwagen et al., 2005) mutants showed weak DSB repair defect with normal CO formation (Fig. 6 B-F). We also checked genomewide DSB repair by examining chromosome bands using pulse-field gel electrophoresis (Fig. S1 E). Different from the *CDC53mn* and *pch2* single mutants, the *CDC53mn pch2* did not recover bands of intact chromosomes at late time points, supporting little DSB repair. These indicate that Pch2 and Cdc53 work distinctly for meiotic DSB repair and/or CO formation. Furthermore, this implies that the completion of the recombination (DSB repair) is not necessary for the formation of full SCs/synapsis. As pointed out by Kleckner (Kleckner et al., 1991), early recombination events are probably sufficient to promote synapsis between homologous chromosomes (Discussion). Consistent with the arrest with unrepaired DSBs, the *CDC53mn pch2* double mutant did not express a pachytene exit marker Cdc5 (Fig. S4 B). The mid-pachytene arrest in the double mutant is induced by recombination checkpoint during meiosis (Hollingsworth and Gaglione, 2019).

**Figure 6.**
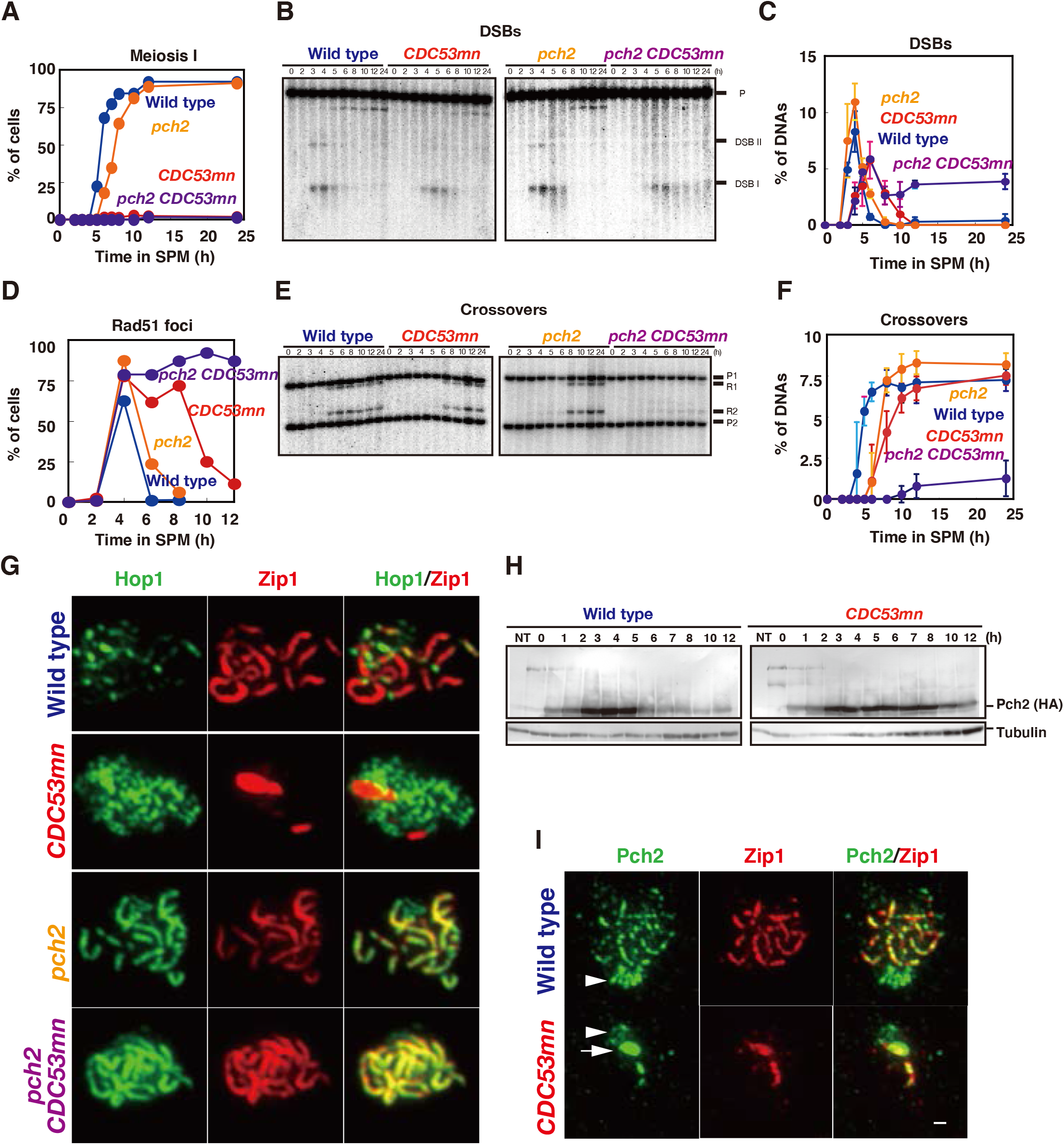
The *pch2 CDC53mn* mutant is defective in meiotic recombination. (A) Entry into meiosis I of various strains was analyzed by DAPI staining. Wild type, NKY1551; *CDC53mn*, ZHY94; *pch2*, ZHY350; *pch2 CDC53mn*, ZHY351. (B, C) DSB repair at the *HIS4-LEU2* locus in various strains was analyzed as described above. Blots are in (B) and quantifications are shown in (C). Error bars were obtained from three independent cultures. (D) Cells containing Rad51 foci were counted at each time point and plotted as described above. Wild type, blue: *CDC53mn*, red; *pch2*, orange; *pch2 CDC53mn*, purple. (E, F) CO formation in various strains was analyzed at the *HIS4-LEU2* locus as described above. Blots are in (E) and quantifications are shown in (F). Error bars were obtained from three independent time courses. (G) Hop1/Zip1 staining in various mutants. Chromosome spreads in each strain were stained with anti-Hop1 (green) and anti-Zip1 (green). (H) Expression of Pch2-HA protein during meiosis. Cell lysate from *PCH2-3HA* and *CDC53mn PCH2-3HA* cells were analyzed by western blotting using anti-HA (top) and anti-tubulin (bottom). (I) Chromosome spreads from various strains were stained with anti-Pch2 (green) and anti-Zip1 (red). Wild type, 4 h; *CDC53mn*, 8 h. Arrows indicate “polycomplex” and arrow heads indicate nucleolus. Bar = 1 μm.

Since Pch2 regulates the localization of Hop1 on chromosomes (Borner et al., 2008), we checked the expression and localization of Hop1 protein (Hollingsworth et al., 1990). In wild type, Hop1 protein was induced after the entry into meiosis and persisted during meiosis (Fig. S4 C). Hop1 showed band shifts from 3 h by the phosphorylation in a manner dependent of Mec1(ATR) kinase (Carballo et al., 2008). These phosphorylated bands of Hop1 decreased during late prophase-I of wild type. The *CDC53mn* mutant induced similar amounts of Hop1 protein as well as phosphorylated Hop1 to the wild type, although the appearance of phospho-Hop1 was delayed by ~2 h due to probably delayed DSB formation (Fig. 2).

We checked the localization of Hop1 by immuno-staining. In wild type, Hop1 shows dotty or short-line staining prior to full synapsis and most of Hop1 dissociates from synaptic regions of pachytene chromosomes (Fig. 6 G). As a result, Hop1 localization of pachytene chromosomes is largely decreased. As reported (Borner et al., 2008), the *pch2* mutant accumulated “unusual” long Hop1 lines during mid-prophase I, which colocalized with Zip1 lines. In the *CDC53mn pch2* mutant, as seen in the *pch2* mutant, Hop1 accumulated on full-length Zip1 lines (Fig. 6 G). These indicate that, unlike Pch2, Cdc53 does not affect Hop1 protein levels and its localization on chromosomes.

One likely possibility to explain the suppression of SC defect in the *CDC53mn* mutant by the *pch2* is that Pch2 is a target of Cdc53-dependnet protein degradation. To check the possibility, we analyzed amounts of Pch2-HA protein by western blotting. Wild-type cells induced Pch2 from 1 h in meiosis. Pch2 peaked at 4 h and disappeared at 6 h (Fig. 6 H). The *CDC53mn* mutant induced Pch2 protein to wild-type levels during meiosis. The amount of Pch2 decreased slightly at late times. This suggests that Pch2 protein level is not affected by Cdc53 depletion. There are little band shifts of Pch2 during meiosis of wild-type and *CDC53mn* mutant strains by modifications (Fig. 6 H).

We checked Pch2 localization using anti-Pch2 antibody. In wild-type cells, as shown previously (San-Segundo and Roeder, 1999), Pch2 localizes to both chromosomes and nucleolus (Fig. 6 I). On the *CDC53mn* spreads, strong signals of Pch2 were seen on Zip1 PCs. This is consistent with the accumulation of Pch2 on Zip1-PC in other synapsis-defective mutants (Herruzo et al., 2016; San-Segundo and Roeder, 1999). The *CDC53mn* cells also showed clustered foci on nucleolus although these Pch2 signal intensities were relatively weaker than those in wild type (arrows in Fig. 6 I). Chromosomal Pch2 signals are much weaker in the *CDC53mn* cells relative to wild type. Since Pch2 localization on chromosomes depends on Zip1 (Herruzo et al., 2016; San-Segundo and Roeder, 1999), reduced loading of Pch2 in the mutant might be due to defective Zip1 loading.

### The *cdc4* mutant shows a defect in SC assembly

We next looked for a F-box protein working with Cdc53 during meiosis. Since *CDC53mn* cells accumulate Cdc6, whose degradation during mitosis depends on Cdc4 (Perkins et al., 2001), we checked a role of Cdc4 in SC formation by using the temperature-sensitive *cdc4-3* mutant in vegetative growth. Previous analysis showed that this mutant accumulated Sic1 at 36 °C with delayed S-phase entry (Sedgwick et al., 2006). The *cdc4-3* mutant showed defective Zip1 elongation at 32 °C, which is a semi-permissive temperature for the mutant, and accumulated Zip1-PCs (Fig. 7 A). Wild-type cells at 32 °C exhibited similar Rad51- and Zip1-staining kinetics to those at 30 °C (compare with Fig. 1) The *cdc4-3* mutant showed normal Rad51-assembly and delayed Rad51-disassembly at 32 °C (Fig. 7 B), implying the role of Cdc4 in meiotic DSB repair. More importantly, the *pch2* deletion also suppressed the Zip1-assembly defect observed in the *cdc4-3* mutant at 32 °C (Fig. 7 C-E). The *cdc4-3 pch2* double mutant formed long Zip1-lines like the *pch2* single mutant (Fig. 7C). The *cdc4-3 pch2* mutant showed little SC disassembly even at late time points (Fig. 7E). Most of the *cdc4-3* mutant cells did not show an arrest at meiosis I and rather shows delayed entry into meiosis I at 32 °C (Fig. 7 F). These results indicate that Cdc4 regulates SC formation together with Cdc53, but not at the onset of anaphase-I.

**Figure 7.**
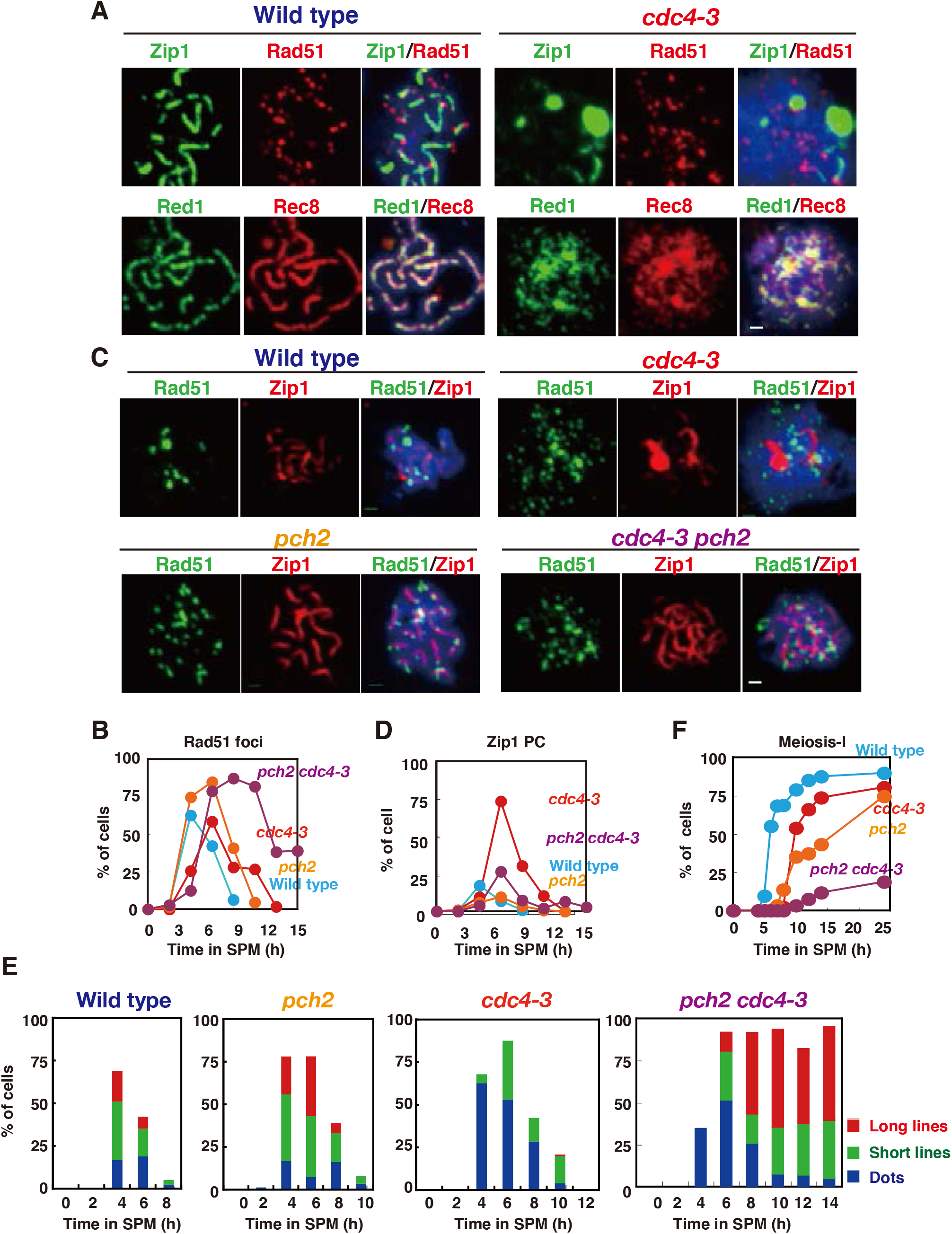
The *cdc4-3* is defective in SC-assembly. (A, B) Chromosome spreads form wild type and *cdc4-3* mutant cells incubated in SPM at 32 °C were stained with anti-Zip1 (green, upper panels) and Rad51 (red, upper panels) and with anti-Red1 (green, bottom panels) and anti-Rec8 (red, bottom panels) as described above. Wild type (NKY1551), 5h; *cdc4-3* (ZHY522) 8 hr. Kinetics of Rad51 focus positive cells are shown in B. Bar = 1 μm. (C-E) Chromosome spreads form various strains at 32 °C were stained with anti-Zip1 (red) and -Rad51 (green). Typical staining patterns were shown in C. Wild type (NKY1551), 5h; *cdc4-3* (ZHY522) 8 h; *pch2* (ZHY350) 8 h; *cdc4-3 pch2* (ZHY580) 8 h. Bar = 1 μm. Kinetics of PC (D) as well as Zip-classes (E) were analyzed as described above. (F) Entry into meiosis I in different strains was analyzed by DAPI staining as described above.

## Discussion

The SCF ubiquitin ligase complexes are involved in various cellular processes during both mitosis and meiosis, particularly in cell cycle progression. In this study, by analyzing the role of two SCF components, Cdc53 (cullin) and Cdc4 (F-box protein) in meiotic cells, we showed that Cdc53 and Cdc4 promote SC formation and that CDC53, but not Cdc4, is required for the transition from the metaphase-I to anaphase-I. Given that Cdc53 as SCF component mediates protein ubiquitylation, it is likely that SCF-dependent ubiquitylation is involved in the two critical meiotic events.

### The role of SCF in SC assembly

The role of ubiquitylation in meiotic chromosome metabolism during meiotic prophase-I has been recently shown by the study of Hei10 ubiquitin ligase in mouse (Qiao et al., 2014). Hei10 plays a direct role in meiotic recombination and an indirect role in SC formation by antagonizing the SUMO ligase Rnf212, a Zip3 ortholog of budding yeast (Qiao et al., 2014). Moreover, in budding yeast, nematode and mouse, the proteasome localizes on meiotic chromosomes (Ahuja et al., 2017; Rao et al., 2017), supporting the role of protein ubiquitylation in meiotic chromosome functions. Our studies showed the role of SCF-dependent ubiquitylation in SC formation.

Like the *zmm* and *ecm11/gmc2* mutants (Humphryes et al., 2013; Pyatnitskaya et al., 2019), the *CDC53mn* mutant is deficient in the SC elongation. Importantly, there are difference in defects in meiotic chromosomal events among these mutants. The *zmm* mutants are defective in CO formation while the *ecm11/gmc2* and *CDC53mn* mutants are proficient. The *ecm11/gmc2* mutant shows normal assembly of ZMM proteins including Zip3 and Msh4. On the other hand, the *CDC53mn* mutant is defective in ZMM assembly except for Msh4-5. These suggest the involvement of Cdc53 in the Msh4/5-independent ZMM function (Pyatnitskaya et al., 2019; Shinohara et al., 2008), which might be unrelated to its role of CO formation.

In *S. cerevisae*, the formation of AEs is coupled with SC elongation accompanied with the polymerization of transverse filaments (Padmore et al., 1991). Moreover, while the *zmm* and *ecm11/gmc2* mutants form normal chromosome axes (Humphryes et al., 2013; Pyatnitskaya et al., 2019), the *CDC53* depletion mutant exhibits altered axis assembly. The SCF^Cdc4^ seems to regulate chromosomal events during early prophase I when axial protein(s) are loaded. *CDC53* depletion seems to cause uncoupling of the loading of SC components, Red1 (and Zip1), with that of Rec8 cohesin and Hop1 to chromosomes. In wild type, the loading of these proteins occurs in early prophase-I in a similar timing. On the other hand, the *CDC53mn* mutant exhibits distinct loading timing between Rec8-Hop1 and Red1-Zip1. Given delay in pre-meiotic S-phase in the *cdc53* mutant, rather than delayed loading of Rec8 and Hop1, promiscuous uncoupled loading of meiosis-specific components such as Red1 (and Zip1) might occur in the mutant. These suggest that coordinated loading of different chromosomal proteins onto chromosomes promotes proper axis formation, which in turn helps SC assembly. We propose that, by functioning in early prophase-I, SCF^Cdc4^ complex controls coordinated loading of axis proteins for proper axis formation, which is critical for coupling of axis assembly with SC elongation.

Although we could not defect any immune-staining signals of Cdc53 on meiosis chromosome spreads, recent studies on mouse spermatocyte showed the localization of a SCF component, Skp1, on the lateral element of SCs (Guan et al., 2020), which supports the role of SCF in axis assembly. It is known that SCF regulates mitotic chromosome condensation in fruit fly and nematode (Buster et al., 2013; Feng et al., 1999). Recently, SCF ubiquitin ligase is shown to promote meiotic chromosome pairing as well as the entry into meiosis in *C. elegans* (Mohammad et al., 2018) and mouse (Gray et al., 2020; Guan et al., 2020), which are shared defects with yeast *CDC53* depletion mutant. Therefore, SCF ubiquitin ligase regulates chromosome morphogenesis during mitotic and meiotic phases in various organisms.

### SC assembly is negatively regulated

Since the SCF complex promotes the ubiquitylation of a target protein for degradation, we could postulate the presence of a negative regulatory pathway for SC assembly, which would be inactivated by the SCF (Fig. 8). In the absence of SCF, the putative negative regulator might inhibit SC elongation. This putative negative regulator might be involved in proper coordination of axis assembly and SC elongation. In this scenario, we could expect to identify a mutation that suppresses the SC defect induced by the *CDC53* depletion. Indeed, we found that the deletion of the *PCH2* gene suppresses a defect in SC assembly in the *CDC53mn* and *cdc4* mutant cells. This suggests the presence of a Pch2-dependent negative regulation for SC assembly. One possibility is that Pch2 could be a direct target of the SCF^Cdc4^-mediated ubiquitylation. However, opposed to this expectation, our studies here did not provide any evidences to support the hypothesis that Cdc53 controls post-translational status of Pch2, Thus, there might be the other target of SCF^Cdc4^ in SC assembly (“X” in Fig. 8).

**Figure 8.**
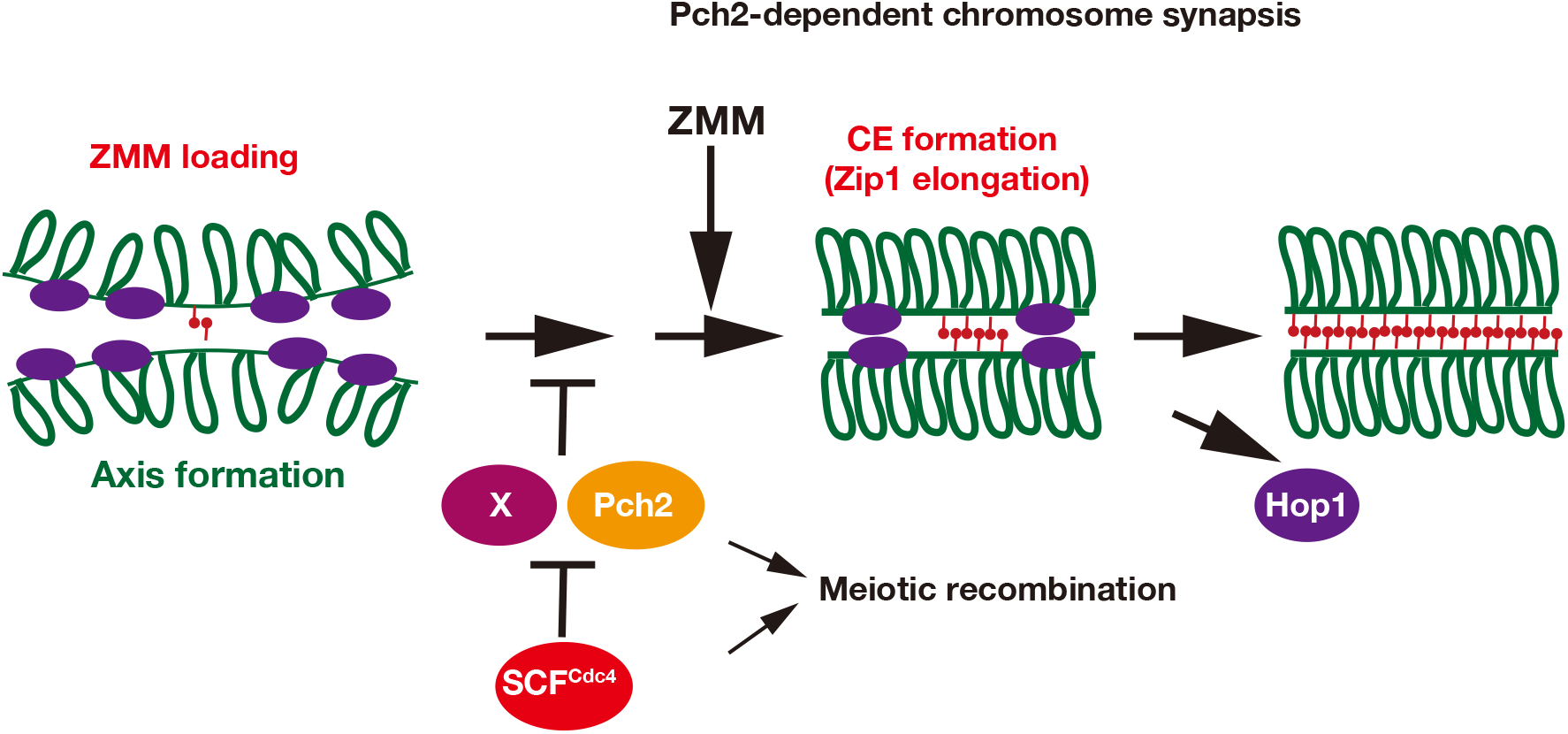
A model on regulation of SC formation by SCF and Pch2. See the text in more detail. The SCF^Cdc4^ promotes SC formation by downregulating Pch2 and protein “X”, both of which may negatively regulate proper axis assembly for SC formation. SCF^Cdc4^ may promote the ubiquitylation of protein “X” for degradation to promote SC assembly.

If Pch2 is not an SCF^Cdc4^ substrate, how does *pch2* deletion suppresses SC-assembly defect in the *CDC53mn* mutant? In SC assembly, Pch2 (TRIP13 in mammals) regulates the dissociation of a chromosome axis protein, Hop1 (HORMAD1/2 in mammals), which might control the synapsis (Borner et al., 2008; Wojtasz et al., 2009). One likely possibility is that SCF^Cdc4^ might control Hop1-mediated regulatory pathway for SC assembly. However, the *CDC53* depletion mutant shows wild-type levels of Hop1 protein with normal phosphorylation as well as normal loading (and unloading) of Hop1 on chromosomes (Fig. 6). Therefore, it is very unlikely that the SCF directly down-regulates Hop1. Rather, we propose that, in the absence of Pch2, cells do not require SCF-dependent control for SC assembly. In other words, the *pch2* mutant cells can form SCs in the presence of the negative regulator for SC assembly (Fig. 8). Pch2 may be able to activate the negative regulator for its action as seen in the activation role of TRIP13 in the spindle assembly checkpoint (Vader, 2015. Alternatively, Pch2 may impose the kinetic barrier for SC formation, which might function in parallel with Cdc53-dependent pathway. The latter role has been proposed to nematode Pch2 ortholog (PCH-2) for chromosome pairing and synapsis (Deshong et al., 2014).

In mouse Skp1 conditional knockdown spermatocytes, chromosome synapsis is partly defective with accumulation of Hormad1/2 proteins on synaptic SC regions, suggesting premature SC disassembly (Guan et al., 2020). In the yeast *CDC53* depletion mutant, SC formation is almost defective with the accumulation of Hop1 protein. The *ndt80* mutation, which induced pachytene arrest, did not fully suppress SC-defect in the *CDC53mn* mutant cells, arguing against premature SC disassembly in the yeast mutant.

### The role of SCF in meiotic recombination

In mouse, SUMO-ligase Rnf212 and ubiquitin ligase Hei10 collaborate in meiotic recombination (Qiao et al., 2014). Budding yeast has a Rnf212 ortholog, Zip3, but does not have a Hei10 ortholog. In this study, we found SCF ubiquitin ligase plays a role not only in SC formation but also a role in meiotic recombination. Rather than cooperation, Zip3 and Cdc53 distinctly control CO formation, since the *zip3 CDC53mn* double mutant is more deficient in CO formation than the *zip3* single mutant. Moreover, Cdc53 is essential for CO formation in the absence of Pch2, which plays a weak role in the recombination in wild-type background (Borner et al., 2008). These suggest a role of Cdc53 in meiotic recombination. Our results support the notion that not only sumolyation, but also ubiquitylation plays a role in recombination during yeast meiosis. This is consistent with the result that deletion of the proteasome component, Pre5, impairs the meiotic recombination (Ahuja et al., 2017).

### Relationship between meiotic recombination and SC formation

Previous studies show intimate relationship or coupling between meiotic recombination and SC formation. It is believed that SC regulates meiotic CO formation (Cahoon and Hawley, 2016; Gao and Colaiacovo, 2018). And also meiotic recombination promotes SC formation (Kleckner, 2006; Padmore et al., 1991). On the other hand, our studies revealed two extreme mutant’s situations, which uncouple meiotic recombination and SC elongation. First, in the case of the *CDC53mn* mutant, meiotic CO forms efficiently in the absence of fully-elongated SCs. This indicates that SC, at least SC elongation, IS NOT required for efficient formation of COs *per se*. We also observed wild-type number of Msh5 foci, which is likely to exhibit non-random distribution like Zip3 foci (Fung et al., 2004; Zhang et al., 2014a; Zhang et al., 2014b), on meiotic chromosomes in the mutant. This suggests normal establishment of ZMM-dependent CO formation may occur in the absence of the *CDC53*, thus SC formation/proper axis assembly, although we do not know the effect of *CDC53* depletion on implementation (and/or maintenance) of CO control. Second, in the *CDC53mn pch2* double mutant, we observed normal SC formation with little CO formation or DSB repair. This suggests that the formation of meiotic recombination products such as COs is not necessary for SC formation. Since SC assembly depends on DSB formation, as suggested previously (Kleckner, 2006), early recombination intermediates such as DSBs and/or single-stranded DNAs at recombination sites are sufficient to trigger SC assembly. Alternatively, like in fruit fly and nematode (Dernburg et al., 1998; McKim and Hayashi-Hagihara, 1998), the *CDC53mn pch2* double mutant may induce minor SC assembly pathway, which is less dependent of the recombination.

### Implication for pachytene checkpoint

The *CDC53mn* mutant, which is defective in SC formation, but is proficient in meiotic recombination, passes through pachytene stage and proceeds at least to metaphase-anaphase I transition. Indeed, the *CDC53mn* mutant expresses Cdc5 and Clb1, which are under the control of Ndt80-dependent mid-pachytene checkpoint. This strongly suggests that abnormal SC in the mutant does not trigger any delay or arrest in meiotic prophase I, suggesting the absence of checkpoint, which monitors synapsis (SC elongation) under the condition. Alternatively, Cdc53 by itself might mediate the synapsis checkpoint signaling.

More interestingly, the *pch2* mutation, when combined with the *CDC53mn*, rather accelerates the cell cycle progression, but induces an arrest, due to inability to repair DSBs, suggesting a role of Pch2 in meiotic recombination, but not in the recombination checkpoint. Consistent with this, in mouse, a mutation of the Pch2 homolog Trip13 is known not to alleviate any defects induced by various meiotic mutations (Li and Schimenti, 2007). In addition, in plant, there is no synaptic checkpoint since meiotic cells progress through cell cycle even in the presence of defective synapsis (Hamant et al., 2006). We propose that SC elongation (synapsis) is not monitored by the surveillance mechanism even in yeast, since meiotic cells has an ability to dismantle abnormal SC with normal CO formation.

### The role of SCF in metaphase-I-to-anaphase-I transition

In addition to roles during prophase-I, SCF might is involved in the transition from metaphase I to anaphase I. The simplest interpretation is that SCF may degrade an inhibitor molecule for APC/C at the transition. The *cdc4* mutant is deficient in SC formation, but is proficient in the transition from metaphase I to anaphase I, suggesting the involvement of a F-box protein other hand Cdc4 in this transition. Similar arrest at metaphase/anaphase-I transition was reported to a yeast mutant of the *RAD6* (Yamashita et al., 2004), which encodes an E2 enzyme for ubiquitylation. The role of Rad6 in SC formation is less clear since the *rad6* mutant also reduces DSB formation, which also affect SC assembly. Moreover, at metaphase/anaphase II transition of *Xenopus* oocytes, SCF is known to promote the degradation of an APC/C inhibitor, Erp1/Emi2 (Nishiyama et al., 2007). We need to identify a target molecule of SCF ubiquitin ligase for not only SC assembly but also the onset of anaphase-I. Indeed, recent two studies showed the role of SCF in the transition in mouse meiosis. One study showed the role of SCF in the activation of Wee1 kinase, which negatively regulates Cyclin-dependent kinase (CDK) in anaphase-I onset (Gray et al., 2020). The other showed that the role of MPF (Cdk) activation in both spermatocytes and oocytes (Guan et al., 2020). The role of SCF in metaphase-I-to-anaphase-I transition seems to be conserved from yeast to higher eukaryotes.

## Materials and methods

### Strains and plasmids

All strains described here are derivatives of SK1 diploids, NKY1551 *(MATα/MAT**a**, lys2/”, ura3/” leu2::hisG/”, his4X-LEU2-URA3/his4B-LEU2, arg4-nsp/arg4-bgl)* except *cdc4-3* strain, which is a congenic strain. An ectopic recombination system with the *URA3-ARG4* cassette was provided by Dr. Michael Lichten. SK1 *cdc4-3* strain and *CENXV-GFP* strains were a kind gift from Drs. D. Stuart and D. Koshland, respectively. The genotypes of each strain used in this study are described in Supplemental Table S1.

### Strain Construction

*pCLB2-3HA-CDC53* were constructed by replacing an endogenous promoter with the promoter from the *CLB2* gene. The addition of the HA tag is important to deplete Cdc53 during meiosis. *pch2, zip3* and *ndt80* null alleles were constructed by PCR-mediated gene disruption using either the *TRP1* or *LEU2* genes (Wach et al., 1994). *REC8-3HA, PCH2-3Flag (−3HA)* and *CDC53-3Flag* were constructed by a PCR-based tagging methodology (De Antoni and Gallwitz, 2000).

### Anti-serum and antibodies

Anti-HA antibody (16B12; Babco), anti-Flag (M2, Sigma), anti-tubulin (MCA77G, Bio-Rad/Serotec, Ltd), anti-GFP (3E6; Molecular Probes), and guinea pig anti-Rad51 (Shinohara et al., 2000) were used for staining. Anti-Cdc53 is a generous gift from Dr. M. Blobel. Anti-Zip1, -Zip3, -Zip2, -Mer3, -Spo22/Zip4, -Msh4, -Msh5 as well as anti-Red1 were described previously (Shinohara et al., 2008). Anti-Rec8 antibody was described previously (Rao et al., 2011). Anti-Hop1 serum was described in (Bani Ismail et al., 2014). Anti-Sic1 (sc-50441, 1:1000) and anti-Cdc5 (sc-33635, 1:1000) antibodies were purchased from SantaCruz Biotech. Anti-Cdc6 (Cdc6 9H8/5) was purchased from Abcam. The second antibodies for staining were Alexa-488 (Goat) and −594 (Goat) IgG used at a 1/2000 dilution (Themo Fishers).

Anti-Pch2 was raised against recombinant N-terminus 300 amino acid of truncated protein purified from *E. coli*. An open-reading frame of the truncated Pch2 was PCR-amplified and inserted into pET15b plasmid (Novagen) in which an N-terminus of the *PCH2* gene was tagged with hexa-histidine. Pch2 protein with the histidine tag was affinity-purified using the Nickel resin as described by manufactures and used for immunization (MBL Co. Ltd).

For double staining the following combinations were used; anti-Rad51 (guinea pig) and anti-Zip1 (rabbit), anti-ZMM (Zip2, Zip3, Zip4/Spo22, Msh4, Msh5, all rabbit) and anti-Zip1 (rat); anti-Red1 (chicken), anti-Rec8 (rabbit); anti-Zip1 (rat), anti-Rec8 (rabbit); anti-Zip1 (rat), anti-Pch2 (rabbit); anti-Zip1 (rat), anti-Hop1 (rabbit).

### Cytology

Immunostaining of chromosome spreads was performed as described previously (Shinohara et al., 2000; Shinohara et al., 2003). Stained samples were observed using an epi-fluorescent microscope (BX51; Olympus) with a 100 X objective (NA1.3). Images were captured by CCD camera (Cool Snap; Roper) and, then processed using IP lab and/or iVision (Sillicon), and Photoshop (Adobe) software. For focus counting, more than 100 nuclei were counted at each time point.

High-resolution images were captured by a computer-assisted fluorescence microscope system (Delta vision, Applied Precision). The objective lens was an oil-immersion lens (100X, NA=1.35). Image deconvolution was carried out using an image workstation (SoftWorks; Applied Precision).

Pairing of chromosomes was analyzed in the whole yeast cells with two homologous LacI-GFP spots at *CENX* locus. Following fluorescence microscope imaging, the number of chromosomal locus-marked GFP foci in a single cell was counted manually.

The distance between two GFP foci on chromosome IV on a single focal plane in a intact yeast cell taken by the fluorescence microscope system (Delta vision, Applied Precision) was measured by NIH image J program.

Zip1 polycomplexes (PCs) were defined as a relatively large Zip1 staining outside of the DAPI staining region.

### Western blotting

Western blotting was performed as described previously (Hayase et al., 2004; Shinohara et al., 2008). Western blotting was performed for cell lysates extracted by TCA method. After being harvested and washed twice with 20% TCA, cells were roughly disrupted by Yasui Kikai (Yasui Kikai Co Ltd, Japan). Protein precipitation recovered by centrifuge at 3000rpm for 5min was suspended in SDS-PAGE sample buffer adjusting to pH8.8 and then boiled for 95°C, 2min.

### Southern Blotting

Time-course analyses of DNA events in meiosis and cell cycle progression were performed as described previously (Shinohara et al., 1997; Storlazzi et al., 1996). Southern blotting analysis was performed with the same procedure as in (Storlazzi et al., 1995). For the *HIS4-LEU2* locus, genomic DNA prepared was digested with *XhoI* (for crossover) and *Psf*I (for meiotic DSBs). For the *URA3-ARG4* locus, the DNA were digested with *Xho*I. Probes for southern blotting were Probe 155 for crossover and Probe 291 for DSB detection on the *HIS4-LEU2* as described in (Xu et al., 1995). Image gauge software (Fujifilm Co. Ltd., Japan) was used for quantification for bands.

### Pulsed field gel electrophoresis

For pulsed field gel electrophoresis (PFGE), chromosomal DNA was prepared in agarose plugs as described in (Bani Ismail et al., 2014; Farmer et al., 2011) and run at 14 °C in a CHEF DR-III apparatus (BioRad) using the field 6V/cm at a 120° angle. Switching times followed a ramp from 15.1 to 25.1 seconds.

### Statistics

Means ± S.D values are shown. Datasets (Fig. 2 K; Fig. 4 F) were compared using the Mann-Whitney U-test (Prism, GraphPad).

## On-line supplemental material

Fig. S1A shows a protein level of Cdc53 during meiosis. Fig.1B-D shows meiotic cell cycle analysis of the *CDC53mn* mutant; S-phase, meiosis I and protein expression. Fig. S1E shows PFGE analysis of meiotic chromosomes for DSB repair. Fig. 2A shows immuno-staining analysis of various ZMM proteins, Zip2, Zip4/Spo22, Mer3 and Msh4in the *CDC53mn* mutant. Fig. S2B reveals the expression of a pachytene exit marker, Cdc5. Fig. S3A-D shows meiotic defects of the *ndt80 CDC53mn* mutant. Fig. S3E shows the localization of Zip1 in the *CDC53mn* and *gcm2* mutants. Fig. S4A, B shows meiotic cell progression analysis of the *CDC53mn pch2* mutant. Fig. S4C is a result of western blotting of Hop1 protein in the *CDC53mn* mutant. Fig. S4D shows images of GFP in the pairing assay for CenXV.

## Acknowledgements

We thank Drs. Neil Hunter and Andreas Hochwagen for discussion. We are grateful for Drs. Mark Goebl, David Stuart, Michael Lichten, and Doug Koshland for materials used in this study. Z.Z. was supported by BMC program and scholarship from the graduate school of science in Osaka university. This work was supported by a Grant-in-Aid from the JSPS KAKENHI Grant Number; 22125001, 22125002, 15H05973 and 16H04742, 19H00981 to A.S. M.S. was supported by the Japanese Society for the Promotion of Science through the Next Generation World-Leading Researchers program (NEXT).

The authors declare no competing financial interest.

## Author contributions

Z.Z., M.S. and A.S. designed the experiments. Z.Z., M.B.I., M.S. and A.S. carried out experiments. M.S provide materials. Z.Z. and A.S. analyzed the data and A.S. wrote the manuscript with inputs from Z.Z. M.B.I. and M.S.

## Abbreviations

CO: crossover
DSB: double-strand breaks
SC: synaptonemal complex
ZMM: Zip-Mer-Msh
SIC: synaptic initiation complex
SCF: Skp-Cullin-F-box

## Supplemental Figures

**Supplemental Figure S1.**
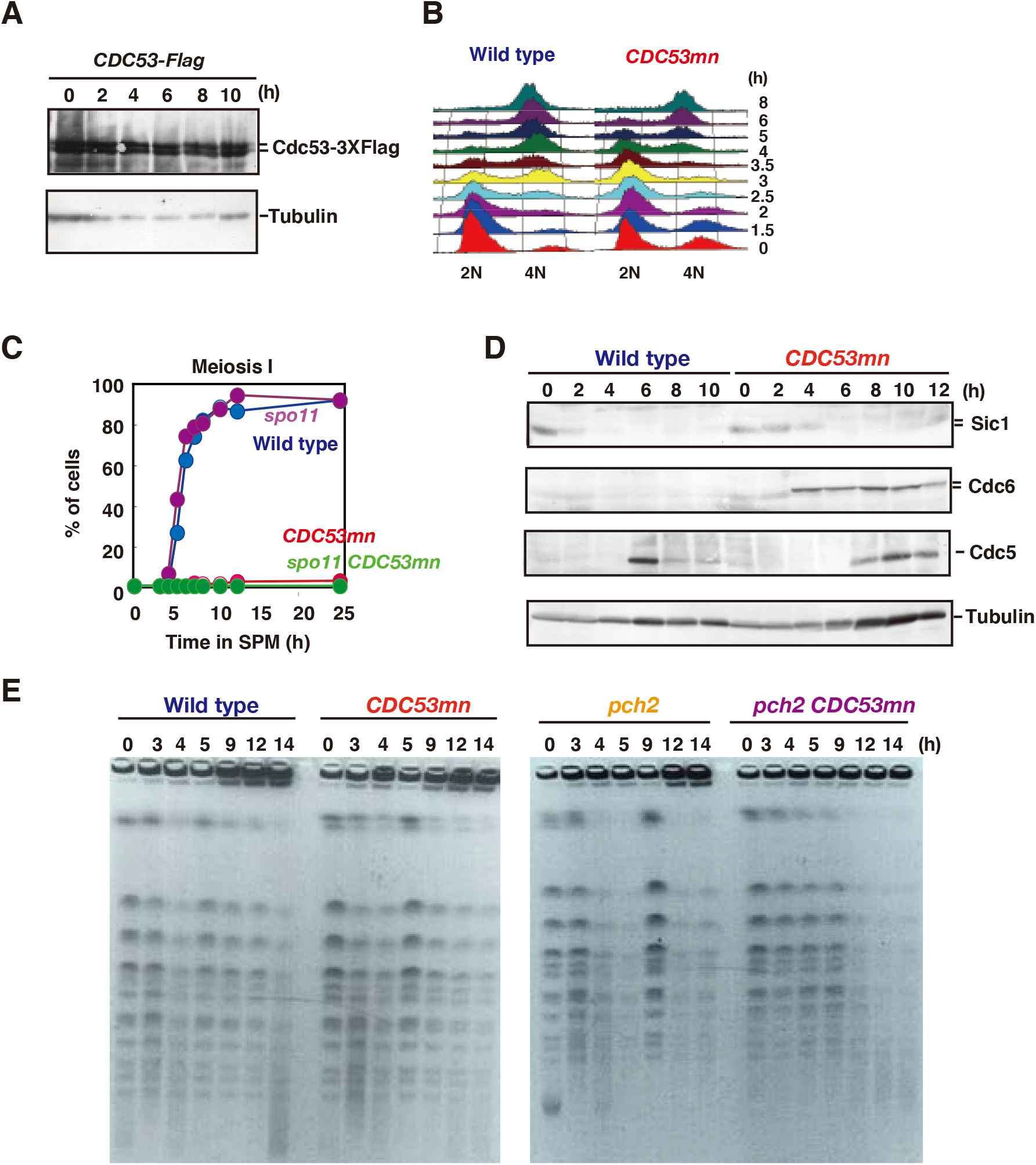
*CDC53mn* mutant shows an arrest during meiosis I. (A) Lysates from meiotic cells with the *CDC53-3XFLAG* gene (ZHY183) were analyzed by western blotting using anti-Flag (upper) and anti-tubulin (lower). (B) Meiotic DNA replication in *CDC53mn*. DNA contents of wild type and *CDC53mn* cells were examined at various times (right) by FACS analysis. (C) Meiosis I progression. Wild type (NKY1551, blue), *CDC53mn* (ZHY94, red), *spo11-Y135F* (MSY1737, purple), *spo11-Y135F CDC53mn* (ZHY272, green) strains were analyzed by DAPI staining. (D) Expression of various proteins in the *CDC53mn* mutant. Lysates obtained from wild type and *CDC53mn* single cells in meiosis were analyzed by Western blotting using anti-Sic1 (upper), anti-Cdc6 (second), anti-Cdc5 (pololike kinase; third) or anti-tubulin (lower) antibodies. (E) CHEF gel analysis of DSB repair during meiosis. Chromosomal DNAs from wild-type (NKY1551) and *CDC53mn* cells (ZHY94), (left panel) and *pch2* (ZHY350) and *pch2 CDC53mn* double mutant (ZHY351) cells were separated by CHEF and gels are stained with ethidium bromide.

**Supplemental Figure S2.**
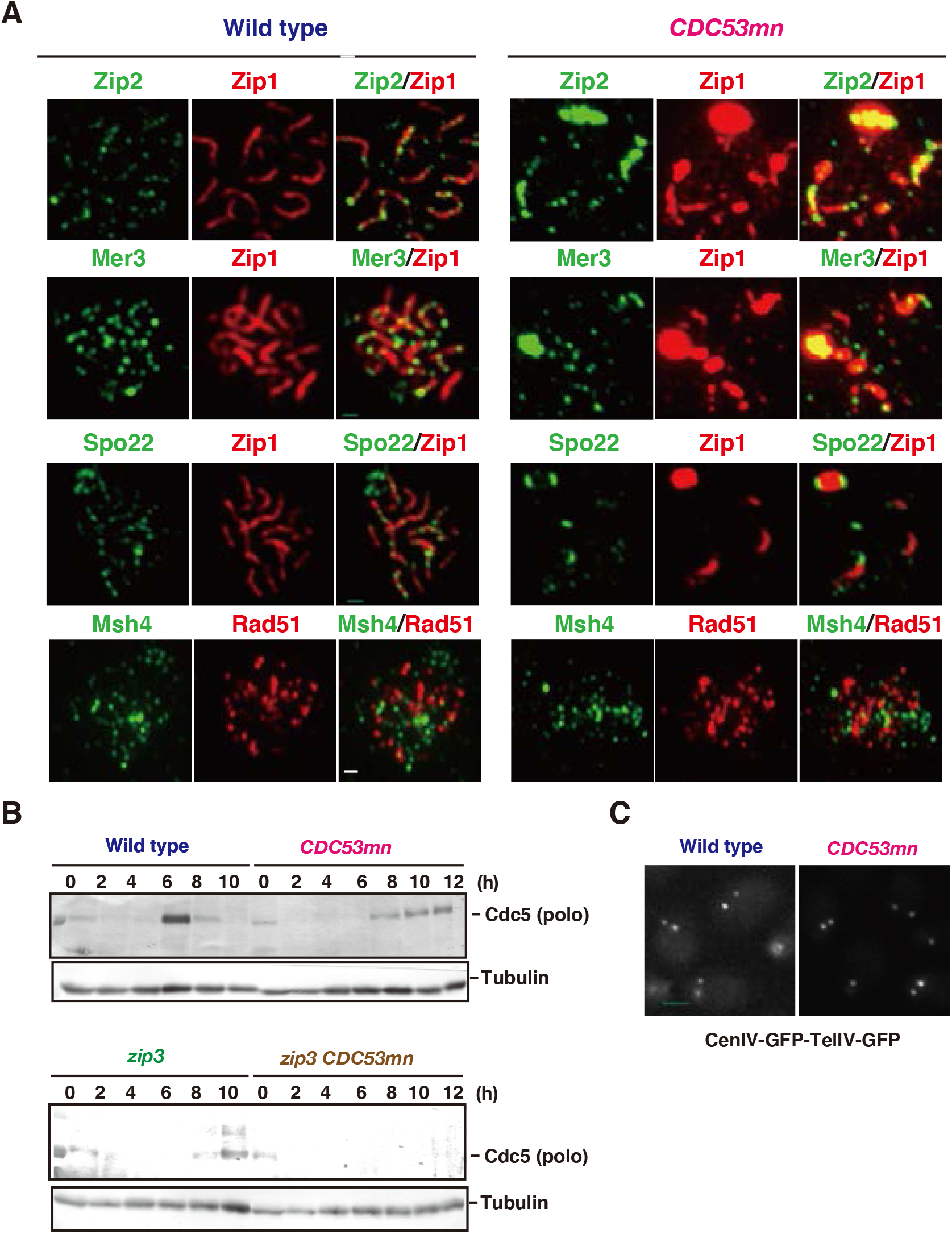
*CDC53* depletion induces abnormal ZMM assembly. (A) Assembly of various ZMM proteins (Zip2, Mer3, Spo22 and Msh4) in the *CDC53mn* mutant. Chromosome spreads from wild type (4 h) and *CDC53mn* mutant (8 h) were stained with anti-Zip2, anti-Mer3, anti -Spo22, or anti-Msh4 antibodies (green) together with anti-Zip1 (red). Wild type, NKY1551; *CDC53mn*, ZHY94. Bar = 1 μm. (B) Western analysis of Cdc5 kinase (top) and tubulin (bottom) in various strains. Wild type, NKY1551; *ndt80*, ZHY516; *CDC53mn*, ZHY94; *ndt80 CDC53mn*, ZHY522.

**Supplemental Figure S3.**
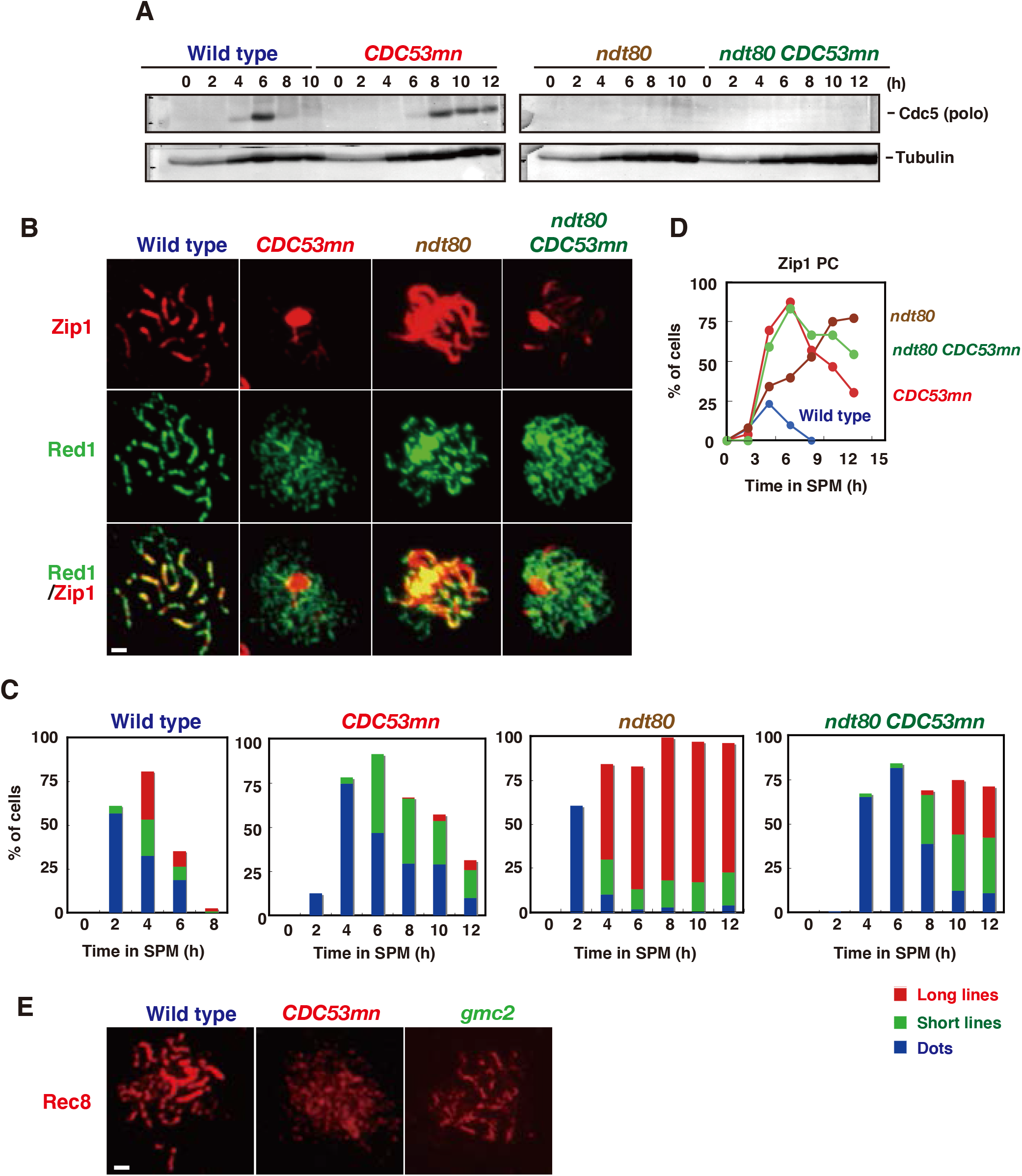
The absence of *NDT80* partially suppresses SC assembly defect in the *CDC53mn*. (A) Expression of Cdc5 polo-like kinase in various strains. Lysates from wild-type, *CDC53mn, ndt80, ndt80 CDC53mn* cells were analyzed by western blotting using anti-Cdc5 (upper) and anti-tubulin (lower). Cdc5 is under the control of Ndt80. Wild type, NKY1551; *ndt80*, ZHY516; *CDC53mn*, ZHY94; *ndt80 CDC53mn*, ZHY522. (B) SC formation in *CDC53mn ndt80* cells. Chromosome spreads from wild-type, *CDC53mn, ndt80, ndt80 CDC53mn* cells were stained with anti-Zip1 (red) and anti-Rec8 (green). Typical images of wild-type (4 h), *CDC53mn* (8 h), *ndt80* (8 h), *ndt80 CDC53mn* (8 h) cells are shown. Bar = 1 μm. (C) Zip1-staining in each strain were classified and plotted at each time point. More than 100 nuclei were counted. Class I (blue bars), Zip1 dots; Class II (green bars), partial Zip1 linear lines; Class III (red bars), linear Zip1 staining. (D) Cells containing Zip1 polycomplexes (PCs) were counted at each time point and plotted. Wild type, blue: *CDC53mn*, red; *ndt80* brown; *ndt80 CDC53mn*, green. (E) Rec8 staining (Red) in wild-type, *CDC53mn* and the *gmc2* mutants.

**Supplemental Figure S4.**
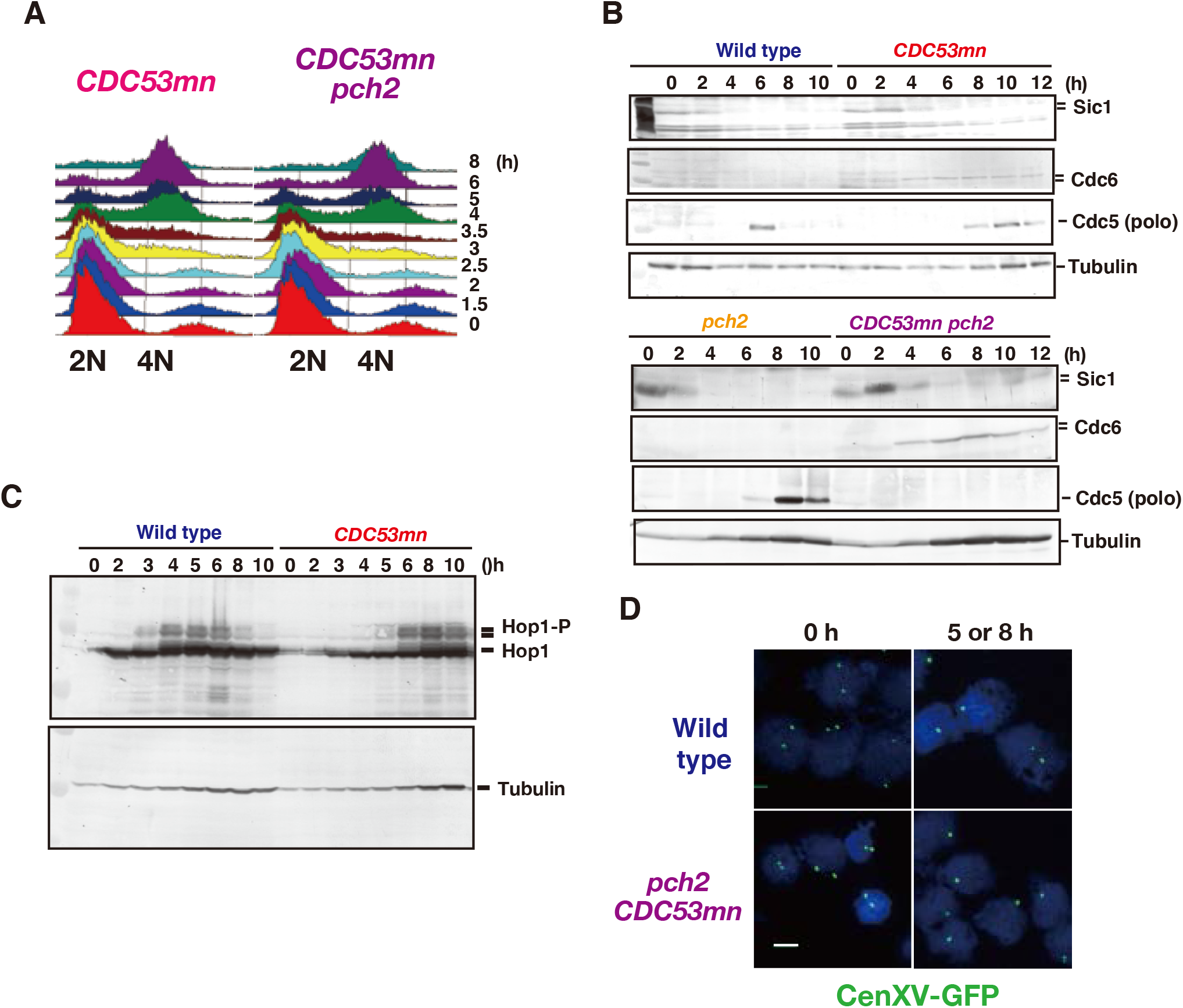
The *pch2 CDC53mn* double mutant arrests at midpachytene stage. (A) Meiotic DNA replication in the *CDC53mn* (ZHY94) and *pch2 CDC53mn* mutants (ZHY351) was analyzed by FACS. (B) Expression of various proteins in the *CDC53mn* mutant. Lysates obtained from wild type and *CDC53mn* single cells in meiosis were analyzed by Western blotting using anti-Sic1 (upper), anti-Cdc6 (second), anti-Cdc5 (polo-like kinase; third) or anti-tubulin (lower) antibodies. (C) Expression of Hop1 in the *CDC53mn* mutant. Lysates obtained from wild type (NKY1551) and *CDC53mn* (ZHY94) cells in meiosis were analyzed by Western blotting using anti-Hop1. (D) Pairing of centromere *XV*. Chromosome spreads from stains homozygous for LacI-GFP-Cen-XV at 0 h and 5 or 8 h were stained with anti-GFP antibody (green) and DAPI (blue) and examined under the epi-fluorescence microscope. Cells with either single or two spots of GFP were counted. More than 100 cells were analyzed at each time point. Percent of cell with single GFP spot is shown in the right. Wild type, ZHY770; *pch2 CDC53mn*, ZHY 772. Bar = 2 μm.

**Table S1.**
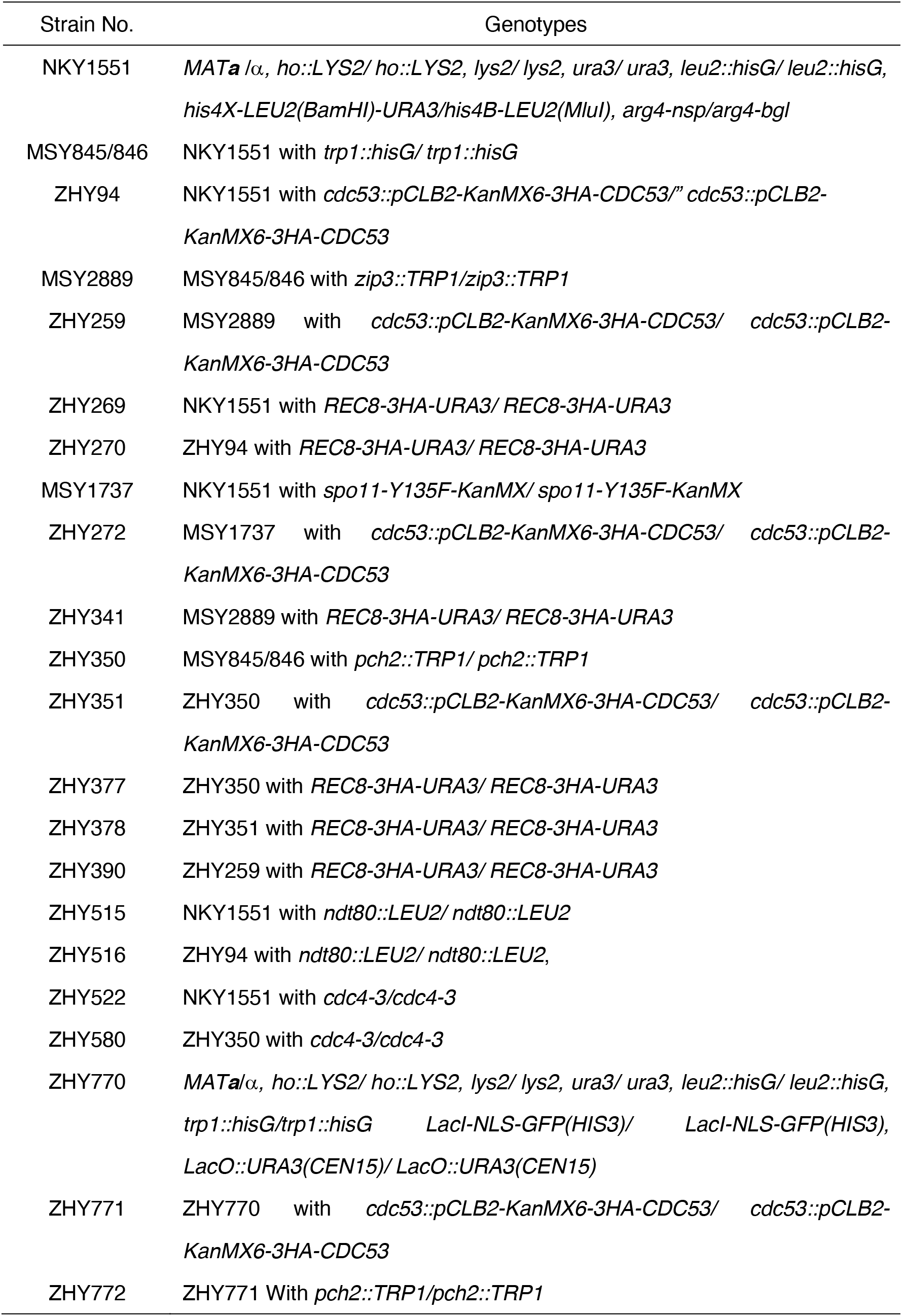
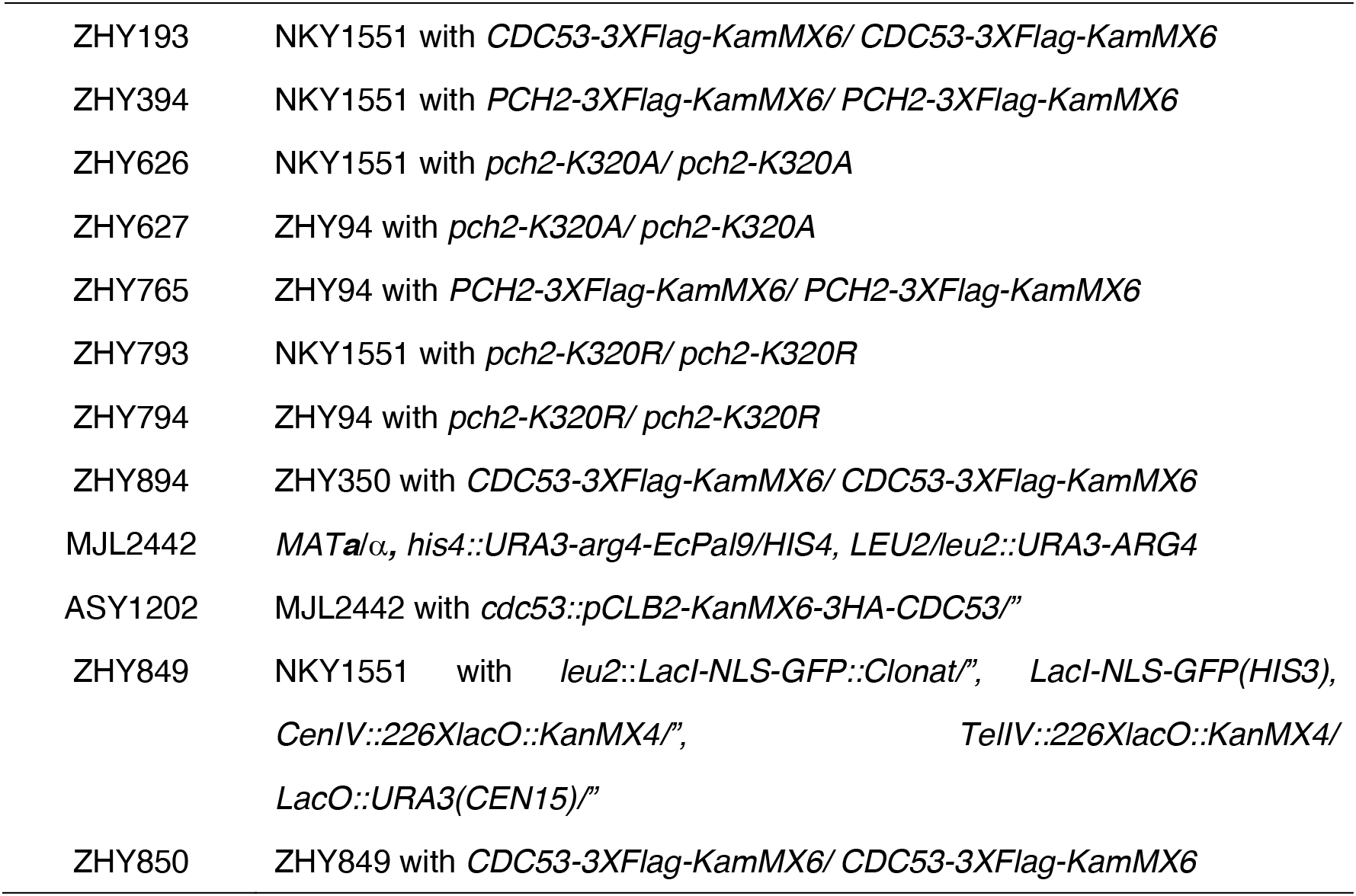
Strain List

| Strain No. | Genotypes |
| --- | --- |
| NKY1551 | *MATa* /$\alpha$, *ho::LYS2/ ho::LYS2, lys2/ lys2, ura3/ ura3, leu2::hisG/ leu2::hisG, his4X-LEU2(BamHI)-URA3/his4B-LEU2(MluI), arg4-nsp/arg4-bgl* |
| MSY845/846 | NKY1551 with *trp1::hisG/ trp1::hisG* |
| ZHY94 | NKY1551 with *cdc53::pCLB2-KanMX6-3HA-CDC53/" cdc53::pCLB2-KanMX6-3HA-CDC53* |
| MSY2889 | MSY845/846 with *zip3::TRP1/zip3::TRP1* |
| ZHY259 | MSY2889 with *cdc53::pCLB2-KanMX6-3HA-CDC53/ cdc53::pCLB2-KanMX6-3HA-CDC53* |
| ZHY269 | NKY1551 with *REC8-3HA-URA3/ REC8-3HA-URA3* |
| ZHY270 | ZHY94 with *REC8-3HA-URA3/ REC8-3HA-URA3* |
| MSY1737 | NKY1551 with *spo11-Y135F-KanMX/ spo11-Y135F-KanMX* |
| ZHY272 | MSY1737 with *cdc53::pCLB2-KanMX6-3HA-CDC53/ cdc53::pCLB2-KanMX6-3HA-CDC53* |
| ZHY341 | MSY2889 with *REC8-3HA-URA3/ REC8-3HA-URA3* |
| ZHY350 | MSY845/846 with *pch2::TRP1/ pch2::TRP1* |
| ZHY351 | ZHY350 with *cdc53::pCLB2-KanMX6-3HA-CDC53/ cdc53::pCLB2-KanMX6-3HA-CDC53* |
| ZHY377 | ZHY350 with *REC8-3HA-URA3/ REC8-3HA-URA3* |
| ZHY378 | ZHY351 with *REC8-3HA-URA3/ REC8-3HA-URA3* |
| ZHY390 | ZHY259 with *REC8-3HA-URA3/ REC8-3HA-URA3* |
| ZHY515 | NKY1551 with *ndt80::LEU2/ ndt80::LEU2* |
| ZHY516 | ZHY94 with *ndt80::LEU2/ ndt80::LEU2,* |
| ZHY522 | NKY1551 with *cdc4-3/cdc4-3* |
| ZHY580 | ZHY350 with *cdc4-3/cdc4-3* |
| ZHY770 | *MATa*/$\alpha$, *ho::LYS2/ ho::LYS2, lys2/ lys2, ura3/ ura3, leu2::hisG/ leu2::hisG, trp1::hisG/trp1::hisG LacI-NLS-GFP(HIS3)/ LacI-NLS-GFP(HIS3), LacO::URA3(CEN15)/ LacO::URA3(CEN15)* |
| ZHY771 | ZHY770 with *cdc53::pCLB2-KanMX6-3HA-CDC53/ cdc53::pCLB2-KanMX6-3HA-CDC53* |
| ZHY772 | ZHY771 With *pch2::TRP1/pch2::TRP1* |
| ZHY193 | NKY1551 with *CDC53-3XFlag-KamMX6/ CDC53-3XFlag-KamMX6* |
| ZHY394 | NKY1551 with *PCH2-3XFlag-KamMX6/ PCH2-3XFlag-KamMX6* |
| ZHY626 | NKY1551 with *pch2-K320A/ pch2-K320A* |
| ZHY627 | ZHY94 with *pch2-K320A/ pch2-K320A* |
| ZHY765 | ZHY94 with *PCH2-3XFlag-KamMX6/ PCH2-3XFlag-KamMX6* |
| ZHY793 | NKY1551 with *pch2-K320R/ pch2-K320R* |
| ZHY794 | ZHY94 with *pch2-K320R/ pch2-K320R* |
| ZHY894 | ZHY350 with *CDC53-3XFlag-KamMX6/ CDC53-3XFlag-KamMX6* |
| MJL2442 | *MATa/α, his4::URA3-arg4-EcPal9/HIS4, LEU2/leu2::URA3-ARG4* |
| ASY1202 | MJL2442 with *cdc53::pCLB2-KanMX6-3HA-CDC53"* |
| ZHY849 | NKY1551 with *leu2::LacI-NLS-GFP::Clonat"*, *LacI-NLS-GFP(HIS3),*<br>*CenIV::226XlacO::KanMX4"*, *TelIV::226XlacO::KanMX4/*<br>*LacO::URA3(CEN15)"/* |
| ZHY850 | ZHY849 with *CDC53-3XFlag-KamMX6/ CDC53-3XFlag-KamMX6* |

## Notes

### Competing Interest Statement

The authors have declared no competing interest.

## References

Agarwal, S., and G.S. Roeder. 2000. Zip3 provides a link between recombination enzymes and synaptonemal complex proteins. Cell. 102:245–255.

Ahuja, J.S., R. Sandhu, R. Mainpal, C. Lawson, H. Henley, P.A. Hunt, J.L. Yanowitz, and G.V. Borner. 2017. Control of meiotic pairing and recombination by chromosomally tethered 26S proteasome. Science (New York, N.Y.). 355:408–411.

Alani, E., R. Padmore, and N. Kleckner. 1990. Analysis of wild-type and *rad50* mutants of yeast suggests an intimate relationship between meiotic chromosome synapsis and recombination. Cell. 61:419–436.

Allers, T., and M. Lichten. 2001. Differential timing and control of noncrossover and crossover recombination during meiosis. Cell. 106:47–57.

Bani Ismail, M., M. Shinohara, and A. Shinohara. 2014. Dot1-dependent histone H3K79 methylation promotes the formation of meiotic double-strand breaks in the absence of histone H3K4 methylation in budding yeast. PloS one. 9:e96648.

Baudat, F., K. Manova, J.P. Yuen, M. Jasin, and S. Keeney. 2000. Chromosome synapsis defects and sexually dimorphic meiotic progression in mice lacking Spo11. Molecular cell. 6:989–998.

Bergerat, A., B. de Massy, D. Gadelle, P.C. Varoutas, A. Nicolas, and P. Forterre. 1997. An atypical topoisomerase II from Archaea with implications for meiotic recombination. Nature. 386:414–417.

Bishop, D.K. 1994. RecA homologs Dmc1 and Rad51 interact to form multiple nuclear complexes prior to meiotic chromosome synapsis. Cell. 79:1081–1092.

Bishop, D.K., D. Park, L. Xu, and N. Kleckner. 1992. *DMC1:* a meiosis-specific yeast homolog of *E. coli recA* required for recombination, synaptonemal complex formation, and cell cycle progression. Cell. 69:439–456.

Borner, G.V., A. Barot, and N. Kleckner. 2008. Yeast Pch2 promotes domainal axis organization, timely recombination progression, and arrest of defective recombinosomes during meiosis. Proceedings of the National Academy of Sciences of the United States of America. 105:3327–3332.

Borner, G.V., N. Kleckner, and N. Hunter. 2004. Crossover/noncrossover differentiation, synaptonemal complex formation, and regulatory surveillance at the leptotene/zygotene transition of meiosis. Cell. 117:29–45.

Buster, D.W., S.G. Daniel, H.Q. Nguyen, S.L. Windler, L.C. Skwarek, M. Peterson, M. Roberts, J.H. Meserve, T. Hartl, J.E. Klebba, D. Bilder, G. Bosco, and G.C. Rogers. 2013. SCFSlimb ubiquitin ligase suppresses condensin II-mediated nuclear reorganization by degrading Cap-H2. The Journal of cell biology. 201:49–63.

Cahoon, C.K., and R.S. Hawley. 2016. Regulating the construction and demolition of the synaptonemal complex. Nature structural & molecular biology. 23:369–377.

Cao, L., E. Alani, and N. Kleckner. 1990. A pathway for generation and processing of double-strand breaks during meiotic recombination in S. cerevisiae. Cell. 61:1089–1101.

Carballo, J.A., A.L. Johnson, S.G. Sedgwick, and R.S. Cha. 2008. Phosphorylation of the axial element protein Hop1 by Mec1/Tel1 ensures meiotic interhomolog recombination. Cell. 132:758–770.

Challa, K., V.G. Fajish, M. Shinohara, F. Klein, S.M. Gasser, and A. Shinohara. 2019. Meiosis-specific prophase-like pathway controls cleavage-independent release of cohesin by Wapl phosphorylation. PLoS genetics. 15:e1007851.

Challa, K., M.S. Lee, M. Shinohara, K.P. Kim, and A. Shinohara. 2016. Rad61/Wpl1 (Wapl), a cohesin regulator, controls chromosome compaction during meiosis. Nucleic acids research. 44:3190–3203.

Chen, C., A. Jomaa, J. Ortega, and E.E. Alani. 2014. Pch2 is a hexameric ring ATPase that remodels the chromosome axis protein Hop1. Proceedings of the National Academy of Sciences of the United States of America. 111:E44–53.

Cheng, C.H., Y.H. Lo, S.S. Liang, S.C. Ti, F.M. Lin, C.H. Yeh, H.Y. Huang, and T.F. Wang. 2006. SUMO modifications control assembly of synaptonemal complex and polycomplex in meiosis of *Saccharomyces cerevisiae*. Genes & development. 20:2067–2081.

Chu, S., and I. Herskowitz. 1998. Gametogenesis in yeast is regulated by a transcriptional cascade dependent on Ndt80. Molecular cell. 1:685–696.

Chua, P.R., and G.S. Roeder. 1998. Zip2, a meiosis-specific protein required for the initiation of chromosome synapsis. Cell. 93:349–359.

Clyne, R.K., V.L. Katis, L. Jessop, K.R. Benjamin, I. Herskowitz, M. Lichten, and K. Nasmyth. 2003. Polo-like kinase Cdc5 promotes chiasmata formation and cosegregation of sister centromeres at meiosis I. Nature cell biology. 5:480–485.

Cooper, K.F., and R. Strich. 2011. Meiotic control of the APC/C: similarities & differences from mitosis. Cell Div. 6:16.

De Antoni, A., and D. Gallwitz. 2000. A novel multi-purpose cassette for repeated integrative epitope tagging of genes in *Saccharomyces cerevisiae*. Gene. 246:179–185.

Dernburg, A.F., K. McDonald, G. Moulder, R. Barstead, M. Dresser, and A.M. Villeneuve. 1998. Meiotic recombination in *C. elegans* initiates by a conserved mechanism and is dispensable for homologous chromosome synapsis. Cell. 94:387–398.

Deshong, A.J., A.L. Ye, P. Lamelza, and N. Bhalla. 2014. A quality control mechanism coordinates meiotic prophase events to promote crossover assurance. PLoS genetics. 10:e1004291.

Eijpe, M., H. Offenberg, R. Jessberger, E. Revenkova, and C. Heyting. 2003. Meiotic cohesin REC8 marks the axial elements of rat synaptonemal complexes before cohesins SMC1beta and SMC3. The Journal of cell biology. 160:657–670.

Farmer, S., W.K. Leung, and H. Tsubouchi. 2011. Characterization of meiotic recombination initiation sites using pulsed-field gel electrophoresis. Methods Mol Biol. 745:33–45.

Feldman, R.M., C.C. Correll, K.B. Kaplan, and R.J. Deshaies. 1997. A complex of Cdc4p, Skp1p, and Cdc53p/cullin catalyzes ubiquitination of the phosphorylated CDK inhibitor Sic1p. Cell. 91:221–230.

Feng, H., W. Zhong, G. Punkosdy, S. Gu, L. Zhou, E.K. Seabolt, and E.T. Kipreos. 1999. CUL-2 is required for the G1-to-S-phase transition and mitotic chromosome condensation in *Caenorhabditis elegans*. Nature cell biology. 1:486–492.

Fung, J.C., B. Rockmill, M. Odell, and G.S. Roeder. 2004. Imposition of crossover interference through the nonrandom distribution of synapsis initiation complexes. Cell. 116:795–802.

Gao, J., and M.P. Colaiacovo. 2018. Zipping and Unzipping: Protein Modifications Regulating Synaptonemal Complex Dynamics. Trends in genetics: TIG. 34:232–245.

Giroux, C.N., M.E. Dresser, and H.F. Tiano. 1989. Genetic control of chromosome synapsis in yeast meiosis. Genome. 31:88–94.

Gray, S., and P.E. Cohen. 2016. Control of Meiotic Crossovers: From Double-Strand Break Formation to Designation. Annual review of genetics. 50:175–210.

Gray, S., E.R. Santiago, J.S. Chappie, and P.E. Cohen. 2020. Cyclin N-Terminal Domain-Containing-1 Coordinates Meiotic Crossover Formation with Cell-Cycle Progression in a Cyclin-Independent Manner. Cell reports. 32:107858.

Guan, Y., N.A. Leu, J. Ma, L. Chmátal, G. Ruthel, J.C. Bloom, M.A. Lampson, J.C. Schimenti, M. Luo, and P.J. Wang. 2020. SKP1 drives the prophase I to metaphase I transition during male meiosis. Science advances. 6:eaaz2129.

Hamant, O., H. Ma, and W.Z. Cande. 2006. Genetics of meiotic prophase I in plants. Annu Rev Plant Biol. 57:267–302.

Hayase, A., M. Takagi, T. Miyazaki, H. Oshiumi, M. Shinohara, and A. Shinohara. 2004. A protein complex containing Mei5 and Sae3 promotes the assembly of the meiosis-specific RecA homolog Dmc1. Cell. 119:927–940.

Herruzo, E., D. Ontoso, S. Gonzalez-Arranz, S. Cavero, A. Lechuga, and P.A. San-Segundo. 2016. The Pch2 AAA+ ATPase promotes phosphorylation of the Hop1 meiotic checkpoint adaptor in response to synaptonemal complex defects. Nucleic acids research. 44:7722–7741.

Hochwagen, A., W.H. Tham, G.A. Brar, and A. Amon. 2005. The FK506 binding protein Fpr3 counteracts protein phosphatase 1 to maintain meiotic recombination checkpoint activity. Cell. 122:861–873.

Hollingsworth, N.M., and R. Gaglione. 2019. The meiotic-specific Mek1 kinase in budding yeast regulates interhomolog recombination and coordinates meiotic progression with double-strand break repair. Current genetics. 65:631–641.

Hollingsworth, N.M., L. Goetsch, and B. Byers. 1990. The *HOP1* gene encodes a meiosis-specific component of yeast chromosomes. Cell. 61:73–84.

Hollingsworth, N.M., L. Ponte, and C. Halsey. 1995. MSH5, a novel MutS homolog, facilitates meiotic reciprocal recombination between homologs in Saccharomyces cerevisiae but not mismatch repair. Genes & development. 9:1728–1739.

Hooker, G.W., and G.S. Roeder. 2006. A Role for SUMO in meiotic chromosome synapsis. Current biology: CB. 16:1238–1243.

Humphryes, N., W.K. Leung, B. Argunhan, Y. Terentyev, M. Dvorackova, and H. Tsubouchi. 2013. The Ecm11-Gmc2 complex promotes synaptonemal complex formation through assembly of transverse filaments in budding yeast. PLoS genetics. 9:e1003194.

Hunter, N. 2015. Meiotic Recombination: The Essence of Heredity. Cold Spring Harbor perspectives in biology. 7.

Kleckner, N. 2006. Chiasma formation: chromatin/axis interplay and the role(s) of the synaptonemal complex. Chromosoma. 115:175–194.

Kleckner, N., R. Padmore, and D.K. Bishop. 1991. Meiotic chromosome metabolism: one view. Cold Spring Harb Symp Quant Biol. 56:729–743.

Klein, F., P. Mahr, M. Galova, S.B. Buonomo, C. Michaelis, K. Nairz, and K. Nasmyth. 1999. A central role for cohesins in sister chromatid cohesion, formation of axial elements, and recombination during yeast meiosis. Cell. 98:91–103.

Koivomagi, M., E. Valk, R. Venta, A. Iofik, M. Lepiku, E.R. Balog, S.M. Rubin, D.O. Morgan, and M. Loog. 2011. Cascades of multisite phosphorylation control Sic1 destruction at the onset of S phase. Nature. 480:128–131.

Lee, B.H., and A. Amon. 2003. Role of Polo-like kinase CDC5 in programming meiosis I chromosome segregation. Science (New York, N.Y.). 300:482–486.

Leung, W.K., N. Humphryes, N. Afshar, B. Argunhan, Y. Terentyev, T. Tsubouchi, and H. Tsubouchi. 2015. The synaptonemal complex is assembled by a polySUMOylation-driven feedback mechanism in yeast. The Journal of cell biology. 211:785–793.

Li, X.C., and J.C. Schimenti. 2007. Mouse pachytene checkpoint 2 (trip13) is required for completing meiotic recombination but not synapsis. PLoS genetics. 3:e130.

Marston, A.L., and A. Amon. 2004. Meiosis: cell-cycle controls shuffle and deal. Nat Rev Mol Cell Biol. 5:983–997.

McKim, K.S., and A. Hayashi-Hagihara. 1998. mei-W68 in Drosophila melanogaster encodes a Spo11 homolog: evidence that the mechanism for initiating meiotic recombination is conserved. Genes & development. 12:2932–2942.

Mohammad, A., K. Vanden Broek, C. Wang, A. Daryabeigi, V. Jantsch, D. Hansen, and T. Schedl. 2018. Initiation of Meiotic Development Is Controlled by Three Post-transcriptional Pathways in Caenorhabditis elegans. Genetics. 209:1197–1224.

Nakagawa, T., and H. Ogawa. 1999. The *Saccharomyces cerevisiae MER3* gene, encoding a novel helicase-like protein, is required for crossover control in meiosis. Embo J. 18:5714–5723.

Nakatsukasa, K., F. Okumura, and T. Kamura. 2015. Proteolytic regulation of metabolic enzymes by E3 ubiquitin ligase complexes: lessons from yeast. Critical reviews in biochemistry and molecular biology. 50:489–502.

Nishiyama, T., K. Ohsumi, and T. Kishimoto. 2007. Phosphorylation of Erp1 by p90rsk is required for cytostatic factor arrest in Xenopus laevis eggs. Nature. 446:1096–1099.

Nottke, A.C., H.M. Kim, and M.P. Colaiacovo. 2017. Wrestling with Chromosomes: The Roles of SUMO During Meiosis. Advances in experimental medicine and biology. 963:185–196.

Novak, J.E., P.B. Ross-Macdonald, and G.S. Roeder. 2001. The budding yeast Msh4 protein functions in chromosome synapsis and the regulation of crossover distribution. Genetics. 158:1013–1025.

Oelschlaegel, T., M. Schwickart, J. Matos, A. Bogdanova, A. Camasses, J. Havlis, A. Shevchenko, and W. Zachariae. 2005. The yeast APC/C subunit Mnd2 prevents premature sister chromatid separation triggered by the meiosis-specific APC/C-Ama1. Cell. 120:773–788.

Okaz, E., O. Arguello-Miranda, A. Bogdanova, P.K. Vinod, J.J. Lipp, Z. Markova, I. Zagoriy, B. Novak, and W. Zachariae. 2012. Meiotic prophase requires proteolysis of M phase regulators mediated by the meiosis-specific APC/CAma1. Cell. 151:603–618.

Padmore, R., L. Cao, and N. Kleckner. 1991. Temporal comparison of recombination and synaptonemal complex formation during meiosis in S. cerevisiae. Cell. 66:1239–1256.

Penkner, A.M., S. Prinz, S. Ferscha, and F. Klein. 2005. Mnd2, an essential antagonist of the anaphase-promoting complex during meiotic prophase. Cell. 120:789–801.

Perkins, G., L.S. Drury, and J.F. Diffley. 2001. Separate SCF(CDC4) recognition elements target Cdc6 for proteolysis in S phase and mitosis. Embo J. 20:4836–4845.

Pesin, J.A., and T.L. Orr-Weaver. 2008. Regulation of APC/C Activators in Mitosis and Meiosis. Annu Rev Cell Dev Biol.

Petronczki, M., M.F. Siomos, and K. Nasmyth. 2003. Un menage a quatre: the molecular biology of chromosome segregation in meiosis. Cell. 112:423–440.

Pyatnitskaya, A., V. Borde, and A. De Muyt. 2019. Crossing and zipping: molecular duties of the ZMM proteins in meiosis. Chromosoma.

Qiao, H., H.B. Prasada Rao, Y. Yang, J.H. Fong, J.M. Cloutier, D.C. Deacon, K.E. Nagel, R.K. Swartz, E. Strong, J.K. Holloway, P.E. Cohen, J. Schimenti, J. Ward, and N. Hunter. 2014. Antagonistic roles of ubiquitin ligase HEI10 and SUMO ligase RNF212 regulate meiotic recombination. Nature genetics. 46:194–199.

Rao, H.B., H. Qiao, S.K. Bhatt, L.R. Bailey, H.D. Tran, S.L. Bourne, W. Qiu, A. Deshpande, A.N. Sharma, C.J. Beebout, R.J. Pezza, and N. Hunter. 2017. A SUMO-ubiquitin relay recruits proteasomes to chromosome axes to regulate meiotic recombination. Science (New York, N.Y.). 355:403–407.

Rao, H.B., M. Shinohara, and A. Shinohara. 2011. Mps3 SUN domain is important for chromosome motion and juxtaposition of homologous chromosomes during meiosis. Genes Cells. 16:1081–1096.

Romanienko, P.J., and R.D. Camerini-Otero. 2000. The mouse Spo11 gene is required for meiotic chromosome synapsis. Molecular cell. 6:975–987.

San-Segundo, P.A., and G.S. Roeder. 1999. Pch2 links chromatin silencing to meiotic checkpoint control. Cell. 97:313–324.

Sedgwick, C., M. Rawluk, J. Decesare, S. Raithatha, J. Wohlschlegel, P. Semchuk, M. Ellison, J. Yates, 3rd, and D. Stuart. 2006. *Saccharomyces cerevisiae* Ime2 phosphorylates Sic1 at multiple PXS/T sites but is insufficient to trigger Sic1 degradation. Biochem J. 399:151–160.

Shinohara, A., H. Ogawa, and T. Ogawa. 1992. Rad51 protein involved in repair and recombination in S. cerevisiae is a RecA-like protein. Cell. 69:457–470.

Shinohara, M., S.L. Gasior, D.K. Bishop, and A. Shinohara. 2000. Tid1/Rdh54 promotes colocalization of Rad51 and Dmc1 during meiotic recombination. Proceedings of the National Academy of Sciences of the United States of America. 97:10814–10819.

Shinohara, M., K. Hayashihara, J.T. Grubb, D.K. Bishop, and A. Shinohara. 2015. DNA damage response clamp 9-1-1 promotes assembly of ZMM proteins for formation of crossovers and synaptonemal complex. Journal of cell science. 128:1494–1506.

Shinohara, M., S.D. Oh, N. Hunter, and A. Shinohara. 2008. Crossover assurance and crossover interference are distinctly regulated by the ZMM proteins during yeast meiosis. Nature genetics. 40:299–309.

Shinohara, M., K. Sakai, T. Ogawa, and A. Shinohara. 2003. The mitotic DNA damage checkpoint proteins Rad17 and Rad24 are required for repair of double-strand breaks during meiosis in yeast. Genetics. 164:855–865.

Shinohara, M., E. Shita-Yamaguchi, J.M. Buerstedde, H. Shinagawa, H. Ogawa, and A. Shinohara. 1997. Characterization of the roles of the *Saccharomyces cerevisiae RAD54* gene and a homologue of *RAD54, RDH54/TID1*, in mitosis and meiosis. Genetics. 147:1545–1556.

Skowyra, D., K.L. Craig, M. Tyers, S.J. Elledge, and J.W. Harper. 1997. F-box proteins are receptors that recruit phosphorylated substrates to the SCF ubiquitin-ligase complex. Cell. 91:209–219.

Smith, A.V., and G.S. Roeder. 1997. The yeast Red1 protein localizes to the cores of meiotic chromosomes. The Journal of cell biology. 136:957–967.

Storlazzi, A., L. Xu, L. Cao, and N. Kleckner. 1995. Crossover and noncrossover recombination during meiosis: timing and pathway relationships. Proc Natl Acad Sci U S A. 92:8512–8516.

Storlazzi, A., L. Xu, A. Schwacha, and N. Kleckner. 1996. Synaptonemal complex (SC) component Zip1 plays a role in meiotic recombination independent of SC polymerization along the chromosomes. Proceedings of the National Academy of Sciences of the United States of America. 93:9043–9048.

Sym, M., J.A. Engebrecht, and G.S. Roeder. 1993. ZIP1 is a synaptonemal complex protein required for meiotic chromosome synapsis. Cell. 72:365–378.

Sym, M., and G.S. Roeder. 1995. Zip1-induced changes in synaptonemal complex structure and polycomplex assembly. The Journal of Cell Biology. 128:455–466.

Tsubouchi, T., H. Zhao, and G.S. Roeder. 2006. The meiosis-specific zip4 protein regulates crossover distribution by promoting synaptonemal complex formation together with Zip2. Dev Cell. 10:809–819.

Voelkel-Meiman, K., L.F. Taylor, P. Mukherjee, N. Humphryes, H. Tsubouchi, and A.J. Macqueen. 2013. SUMO localizes to the central element of synaptonemal complex and is required for the full synapsis of meiotic chromosomes in budding yeast. PLoS genetics. 9:e1003837.

Wach, A., A. Brachat, R. Pohlmann, and P. Philippsen. 1994. New heterologous modules for classical or PCR-based gene disruptions in Saccharomyces cerevisiae. Yeast. 10:1793–1808.

West, A.M., S.C. Rosenberg, S.N. Ur, M.K. Lehmer, Q. Ye, G. Hagemann, I. Caballero, I. Uson, A.J. MacQueen, F. Herzog, and K.D. Corbett. 2019. A conserved filamentous assembly underlies the structure of the meiotic chromosome axis. eLife. 8.

Willems, A.R., T. Goh, L. Taylor, I. Chernushevich, A. Shevchenko, and M. Tyers. 1999. SCF ubiquitin protein ligases and phosphorylation-dependent proteolysis. Philos Trans R Soc Lond B Biol Sci. 354:1533–1550.

Willems, A.R., M. Schwab, and M. Tyers. 2004. A hitchhiker’s guide to the cullin ubiquitin ligases: SCF and its kin. Biochimica et biophysica acta. 1695:133–170.

Wojtasz, L., K. Daniel, I. Roig, E. Bolcun-Filas, H. Xu, V. Boonsanay, C.R. Eckmann, H.J. Cooke, M. Jasin, S. Keeney, M.J. McKay, and A. Toth. 2009. Mouse HORMAD1 and HORMAD2, two conserved meiotic chromosomal proteins, are depleted from synapsed chromosome axes with the help of TRIP13 AAA-ATPase. PLoS genetics. 5:e1000702.

Wu, H.Y., and S.M. Burgess. 2006. Two distinct surveillance mechanisms monitor meiotic chromosome metabolism in budding yeast. Current biology: CB. 16:2473–2479.

Xu, L., M. Ajimura, R. Padmore, C. Klein, and N. Kleckner. 1995. *NDT80*, a meiosis-specific gene required for exit from pachytene in *Saccharomyces cerevisiae*. Mol Cell Biol. 15:6572–6581.

Yamashita, K., M. Shinohara, and A. Shinohara. 2004. Rad6-Bre1-mediated histone H2B ubiquitylation modulates the formation of double-strand breaks during meiosis. Proceedings of the National Academy of Sciences of the United States of America. 101:11380–11385.

Yu, H., J.M. Peters, R.W. King, A.M. Page, P. Hieter, and M.W. Kirschner. 1998. Identification of a cullin homology region in a subunit of the anaphase-promoting complex. Science (New York, N.Y.). 279:1219–1222.

Zachariae, W., A. Shevchenko, P.D. Andrews, R. Ciosk, M. Galova, M.J. Stark, M. Mann, and K. Nasmyth. 1998. Mass spectrometric analysis of the anaphase-promoting complex from yeast: identification of a subunit related to cullins. Science (New York, N.Y.). 279:1216–1219.

Zhang, L., Z. Liang, J. Hutchinson, and N. Kleckner. 2014a. Crossover patterning by the beam-film model: analysis and implications. PLoS genetics. 10:e1004042.

Zhang, L., S. Wang, S. Yin, S. Hong, K.P. Kim, and N. Kleckner. 2014b. Topoisomerase II mediates meiotic crossover interference. Nature. 511:551–556.

Zickler, D., and N. Kleckner. 1999. Meiotic chromosomes: integrating structure and function. Annual review of genetics. 33:603–754.

